# DiffGSP: reversing mRNA diffusion to unlock high-fidelity spatial transcriptomics

**DOI:** 10.64898/2026.09.10.750095

**Authors:** Jixin Liu, Shuli Sun, Yang Xu, Shuai Jiang, Shuhuan Cao, Guojun Li, Xin Zhao, Bingqiang Liu

## Abstract

Spatial transcriptomics enables gene expression profiling within intact tissues while preserving spatial context, providing unprecedented insights into cellular organization and function. However, mRNA diffusion during tissue processing can cause transcripts originating from adjacent cells to be captured at a given spot. This spatial misalignment challenges a fundamental premise of spatial transcriptomics that measured gene expression faithfully corresponds to its spatial origin, thereby compromising spatial fidelity and potentially biasing biological interpretation. Here, we present DiffGSP, a physics-informed framework that integrates Fick’s law with graph signal processing to explicitly model diffusion-induced distortions and recover the underlying spatial gene expression landscape. Comprehensive benchmarking across diverse datasets and evaluation metrics demonstrates the consistent ability of DiffGSP to restore spatial gene expression patterns. By computationally reversing diffusion-induced distortions, DiffGSP enables the discovery of fine anatomical structures in the mouse brain, spatially organized gene modules in the kidney, intratumoral heterogeneity in colorectal cancer, and tertiary lymphoid structures in lung adenocarcinoma. Built on a physically interpretable framework and broadly applicable to sequencing-based spatial transcriptomics technologies, DiffGSP improves the fidelity of spatial gene expression reconstruction and enables more reliable biological interpretation.

## Introduction

A key feature of spatial transcriptomics (ST) is the ability to profile gene expression while preserving spatial information, providing unprecedented opportunities to investigate cellular organization and tissue architecture^1–3^. Among ST technologies, sequencing-based platforms, such as Visium, Slide-seq^4^ and Stereo-seq^5^, stand out for large-scale studies across diverse biological contexts due to their whole-transcriptome profiling capabilities and high scalability^6–8^. In the standard workflow of sequencing-based ST, tissue sections are laid onto capture arrays, and enzymatic permeabilization in the aqueous environment releases cellular mRNAs, which are subsequently captured by the underlying spots^9–11^. These captured mRNAs are then sequenced to determine their identities and reconstruct gene expression profiles of spots.

Due to Brownian diffusion, liberated mRNAs can spread beyond their original locations into neighboring regions, disrupting the correspondence between measured expression signals and spatial locations^12^. Consequently, the expression signal observed at a given location may no longer faithfully represent its true spatial origin, introducing spatial misalignment into ST measurements. This phenomenon compromises the spatial fidelity of ST data, as distorted gene expression profiles propagate into downstream inaccuracies^13^, such as mis-annotating cell types and mis-delimiting spatial domains. Recently, You et al. also underscored mRNA diffusion as a critical challenge in sequencing-based spatial transcriptomics technologies^12^, highlighting its capacity to erode spatial boundaries, misplace gene expression signals, and ultimately distort the interpretation of tissue structure. In SpotClean’s study^14^, contiguously arrayed human and mouse tissues on a single chip during sample preparation revealed rampant cross-species contamination: human transcripts populated ostensibly mouse exclusive spots and vice versa, undermining the credibility of downstream analyses. Such diffusion artifacts are inherent to sequencing-based spatial transcriptomics workflows and remain difficult to mitigate experimentally^15^, perpetuating a critical barrier to accurate biological discovery.

While several computational tools have been developed to improve spatial transcriptomics data quality, such as Sprod^16^, MIST^17^ and spRefine^18^, they primarily focus on denoising rather than addressing the issue of molecular diffusion. Moreover, many of these methods rely on smoothing-based denoising strategies, which may exacerbate diffusion effects^19–21^. Other methods, such as SpotGF^22^, identify genes affected by diffusion and exclude them from downstream analysis, but do not explicitly model or correct the underlying diffusion process.

To date, only a limited number of computational approaches directly address molecular diffusion in spatial transcriptomics data. Among them, SpotClean^14^ employs a statistical framework to distinguish mRNAs originating from within tissue from those diffusing from external sources; however, it relies on strong statistical assumptions rather than physical principles, limiting its ability to faithfully capture the underlying biophysical diffusion process and reducing its cross-platform portability. In addition, the computational complexity of SpotClean constrains its scalability and limits its applicability to large-scale datasets. Beyond molecular diffusion, technical noise further degrades data quality and hampers reliable biological inference. These limitations highlight the need for computational frameworks that explicitly model diffusion physics while maintaining scalability, aiming to restore data fidelity and improve the accuracy and robustness of downstream analyses.

Here, we present DiffGSP, a physics-informed approach that integrates Fick’s diffusion law with graph signal processing (GSP)^23–26^ to correct diffusion-induced distortion in sequencing-based spatial transcriptomics. Unlike previous methods that rely heavily on rigid statistical assumptions, DiffGSP grounds itself in Fick’s law, explicitly modeling the biophysical process of molecular diffusion and providing a physically interpretable framework that improves generalizability across diverse spatial transcriptomics platforms^27, 28^. The inverse diffusion problem is inherently ill-posed and unstable, requiring careful regularization for reliable recovery^29^. To address this challenge, we formulate spatial transcriptomic recovery as an inverse diffusion problem on a spatial graph and develop a graph spectral reconstruction strategy to stabilize the ill-posed inversion. This formulation enables the integration of inverse diffusion solving with graph-based spectral filtering^30–32^ as a regularization strategy for stabilizing the inverse solution. Specifically, DiffGSP integrates inverse diffusion reconstruction with spectral filtering^33^ across both spatial and gene domains in a unified graph spectral framework, enabling joint recovery of pre-diffusion expression signals and suppression of technical noise, thereby improving stability. Together, DiffGSP provides a physics-informed graph spectral framework for stable reconstruction of spatial gene expression signals and recovery of high-quality spatial transcriptomic profiles.

We systematically benchmarked DiffGSP against existing methods, demonstrating its scalability and improved performance in recovering spatial gene expression patterns across diverse datasets and evaluation metrics. We further demonstrated the utility of DiffGSP across diverse biological contexts, ranging from normal tissues to tumor ecosystems, by analyzing datasets generated from multiple spatial transcriptomics platforms, such as ST, Visium, Slide-seqV2, Stereo-seq, and Visium HD. These results highlight the robustness and versatility of DiffGSP across diverse spatial resolutions, sequencing technologies, and experimental settings. Applying DiffGSP to diverse biological systems, we recovered fine anatomical structures in the mouse brain, identified spatially organized gene modules in the kidney, revealed intratumoral heterogeneity in colorectal cancer, and detected tertiary lymphoid structures in lung adenocarcinoma.

Overall, DiffGSP provides a physics-informed framework for modeling and correcting mRNA diffusion effects in spatial transcriptomics, enabling recovery of high-fidelity spatial expression patterns and more reliable downstream biological interpretation across diverse tissues and platforms. Built on a rigorous theoretical foundation and broadly compatible with sequencing-based spatial transcriptomics technologies, DiffGSP serves as a powerful framework for improving spatial transcriptomic analysis and facilitating biological discovery.

## Results

### Overview of DiffGSP

Ideally, each spot would capture transcripts exclusively from its corresponding tissue location. However, molecular diffusion causes transcripts captured at a spot to include contributions from surrounding tissue regions, thereby obscuring true gene expression patterns (**Fig. 1a**). This diffusion-induced signal contamination is commonly observed across diverse tissue samples and sequencing-based spatial transcriptomics platforms (**Supplementary Figs. 1–4**). This contamination adversely affects downstream analyses, compromising multiple downstream tasks, ultimately affecting the accuracy and reliability of biological interpretation. To address this challenge, we developed DiffGSP, a physics-informed graph signal processing framework that explicitly models molecular diffusion and reconstructs pre-diffusion spatial gene expression profiles. DiffGSP formulates mRNA diffusion according to Fick’s law as a partial differential equation (PDE) and represents the diffusion process on a spatial graph using graph signal processing. By transforming diffusion correction into a graph-based inverse diffusion problem, DiffGSP leverages graph spectral regularization to stabilize the ill-posed inversion and recover diffusion-distorted spatial expression landscapes.

**Fig. 1.**
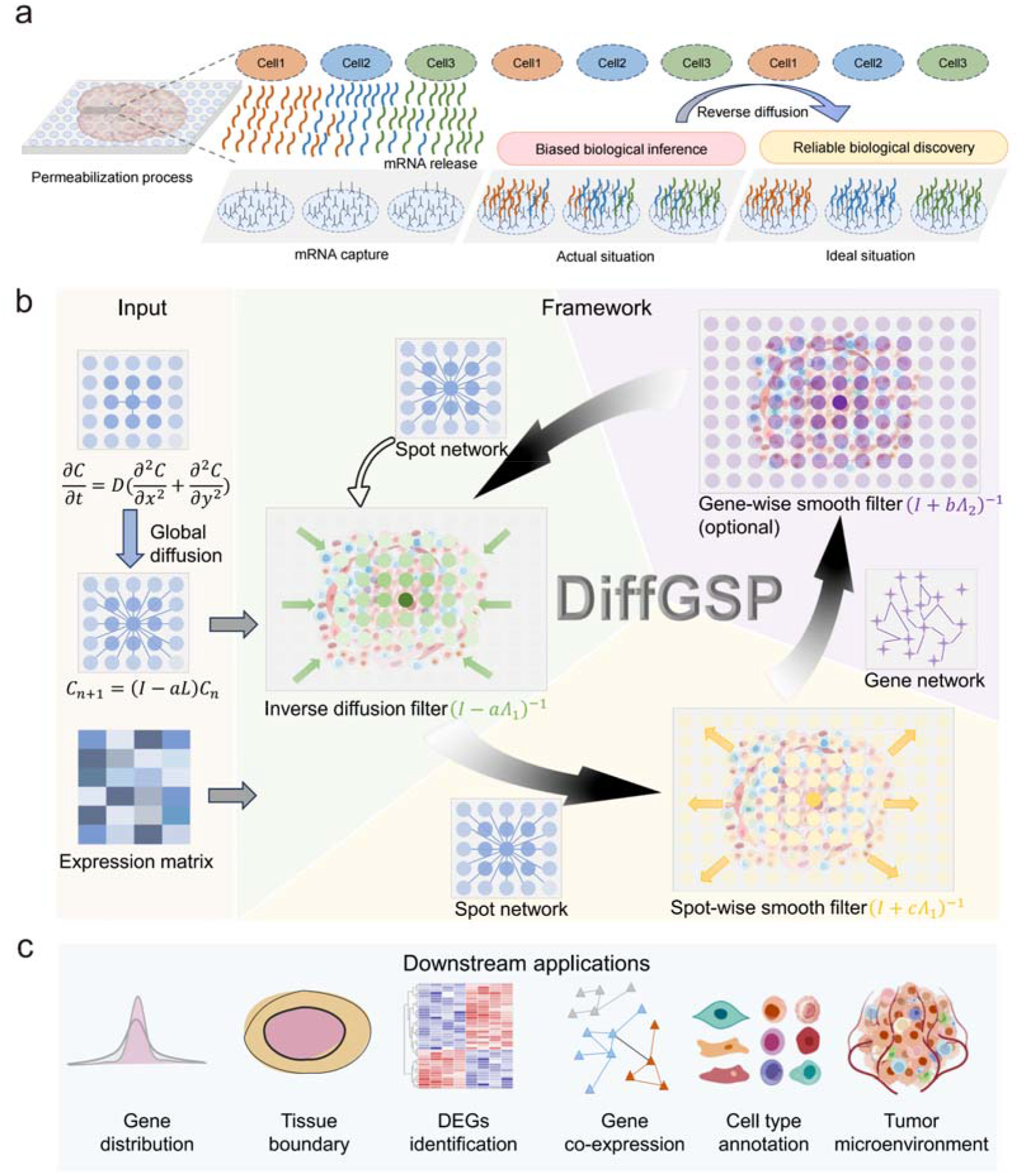
mRNA diffusion and the framework of DiffGSP. **a**, Illustration of mRNA diffusion in sequencing-based spatial transcriptomics. Ideally, transcripts are captured by the spots directly beneath their source cells. However, diffusion can cause mRNAs to spread into neighboring or even distant spots, compromising the accuracy of downstream analyses. **b**, Framework of the DiffGSP algorithm. The process includes inverse diffusion filtering, spot-wise smoothing, gene-wise smoothing, and final output generation. **c**, The downstream analysis tasks of DiffGSP. DiffGSP aims to improve data quality and it can be used in multiple downstream scenarios.

Briefly, DiffGSP constructs a spatial neighboring graph to represent spatial locations and their relationships (**Fig. 1b**). Within this framework, the expression profile of each gene is represented as a graph signal defined over spatial locations. The diffusion process is modeled as signal propagation over the spatial graph, capturing the spatial spreading of molecular signals and their evolution over time. Because diffusion is inherently a forward process that progressively smooths spatial signals, recovering the underlying expression state prior to diffusion requires reversing this process, which introduces numerical instability and requires carefully designed strategies for stable reconstruction. By formulating diffusion correction within a graph signal processing framework, DiffGSP provides a physically interpretable reconstruction strategy without relying on strong statistical assumptions or deep learning models. Furthermore, the graph-based formulation enables broad applicability across diverse tissues and sequencing-based spatial transcriptomics platforms with varying spatial resolutions.

To stabilize inverse diffusion reconstruction, DiffGSP incorporates graph-based low-pass spectral filtering and adaptive diffusion coefficient selection as complementary strategies (**Fig. 1b**). In the graph spectral domain, inverse diffusion can amplify high-frequency components that are sensitive to perturbations, resulting in unstable signal amplification and numerical instability. DiffGSP therefore introduces low-pass spectral filtering as a regularization strategy to constrain these unstable components while preserving smooth and biologically meaningful spatial expression patterns^23, 34, 35^. Specifically, DiffGSP considers two complementary graph structures: a spot-level spatial graph capturing spatial relationships among locations and a gene-level graph representing gene co-expression relationships. Spectral regularization on these graphs enables the integration of spatial and molecular constraints during reconstruction. Inspired by Taubin’s algorithm^36^, DiffGSP combines inverse diffusion reconstruction with graph spectral filtering within a unified framework. In addition, the corresponding parameters can be adaptively estimated when out-of-tissue spots are available by minimizing reconstructed signals in background regions using the L-BFGS-B optimization algorithm^37, 38^. Because excessive diffusion correction can induce unstable amplification during inverse reconstruction, whereas insufficient correction may fail to recover diffusion-induced distortions, DiffGSP further applies the Kneedle algorithm^39^ to identify the transition point across candidate diffusion coefficients. Further details on diffusion coefficient estimation and stabilization strategies are provided in the Methods section. This adaptive strategy balances diffusion correction and numerical stability, enabling reliable recovery of pre-diffusion spatial expression profiles.

By recovering diffusion-distorted spatial expression profiles, DiffGSP substantially enhances downstream spatial transcriptomics analyses (**Fig. 1c**), enabling sharper visualization of spatial expression patterns, more accurate delineation of tissue boundaries, improved detection of differentially expressed genes (DEGs), refined gene co-expression network reconstruction, enhanced cell-type annotation, and more precise characterization of complex tissue microenvironments. Together, these improvements restore spatial fidelity lost during transcript capture and enable more accurate reconstruction of tissue organization, cellular interactions, and spatially resolved biological processes.

In summary, DiffGSP establishes a principled framework for addressing molecular diffusion, an inherent challenge in spatial transcriptomics, by integrating physics-informed inverse diffusion modeling with graph signal processing. By restoring high-fidelity spatial expression landscapes and recovering biologically meaningful patterns, DiffGSP expands the analytical capabilities of spatial transcriptomics and enables more reliable biological discovery.

### DiffGSP reverses molecular diffusion to recover high-fidelity spatial transcriptomic landscapes

To directly evaluate the ability of DiffGSP to correct molecular diffusion, we first analyzed three human–mouse chimeric tissue sections^14^ (S1–S3) profiled using the Visium platform, where adjacent human and mouse tissues provide an intrinsic benchmark for assessing diffusion-induced transcript contamination. Human and mouse tissues were placed contiguously on the same capture array and annotated based on H&E images (**Fig. 2a**). DiffGSP and SpotClean were separately applied to human- and mouse-derived transcript profiles, with annotated tissue regions defined as in-tissue spots and the remaining regions as out-of-tissue spots. We quantified spatial specificity using the log-fold change (LogFC) of mean normalized expression between in-tissue and out-of-tissue spots, where higher values indicate stronger spatial confinement of species-specific transcripts. Both DiffGSP and SpotClean increased species specificity compared with the original data, confirming the presence of diffusion-induced cross-species contamination. Notably, DiffGSP achieved higher spatial specificity than SpotClean, demonstrating its superior ability to restore spatial fidelity compromised by molecular diffusion (**Fig. 2b**). In section S2, DiffGSP produced cleaner species-specific spatial patterns that more closely matched manual tissue annotations (**Fig. 2c**), with consistent improvements observed in sections S1 and S3 (**Supplementary Fig. 5a, c**).

**Fig. 2.**
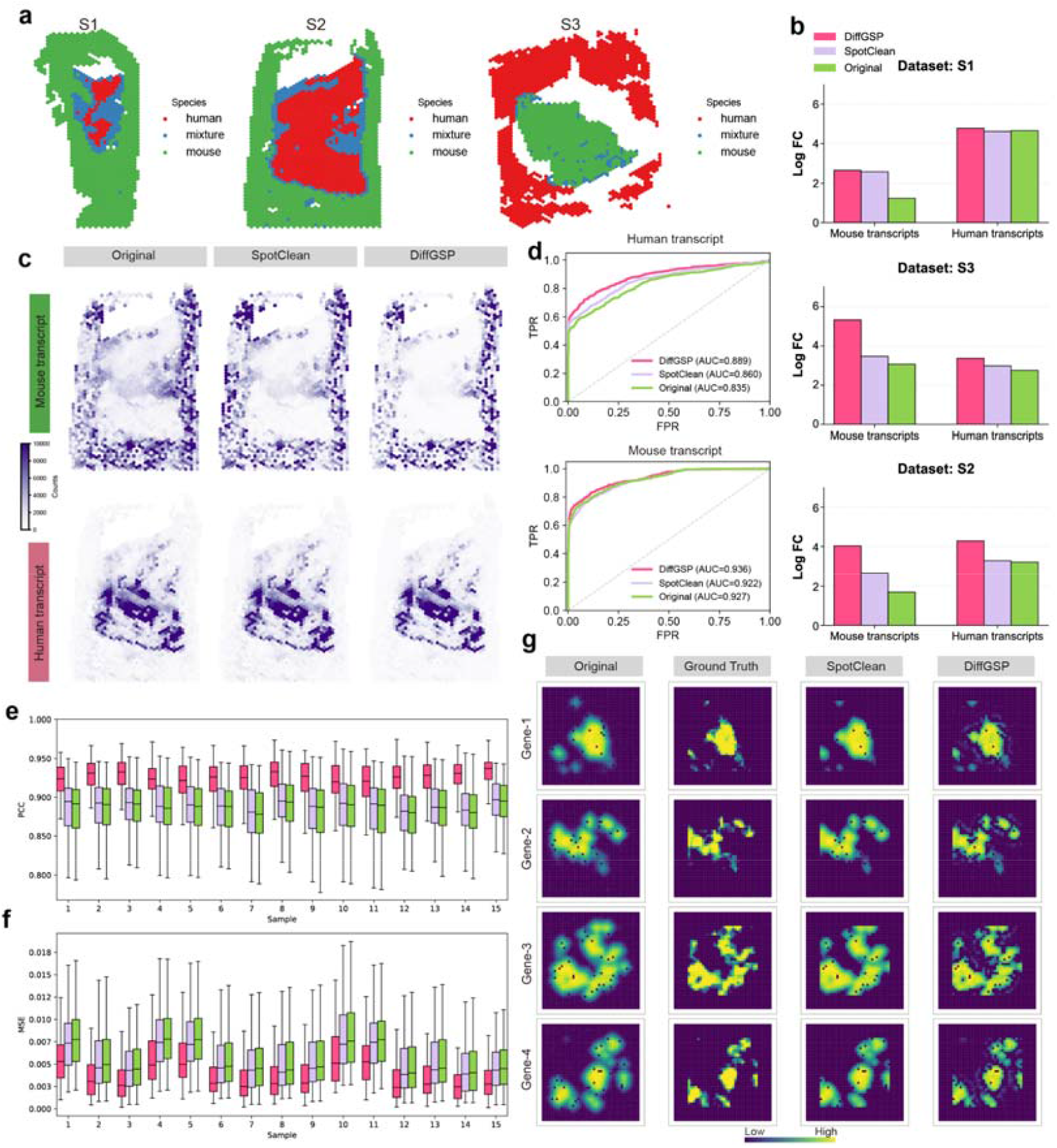
DiffGSP recovers gene distributions and reduces experimental artifacts. **a**, Spatial layouts of three human–mouse chimeric tissue sections (S1, S2, and S3) labeled with species-specific spot annotations (human, mixture, or mouse). **b**, Comparison of Log Fold Change (LogFC) values across three datasets (S1, S3, and S2) processed by different methods: Original (green), SpotClean (purple), and DiffGSP (pink). For human transcripts, LogFC is calculated as the log-transformed ratio of the mean human transcript count in human-associated tissue spots to that in spots outside the human tissue region. For mouse transcripts, LogFC is calculated analogously using mouse-associated tissue spots and spots outside the mouse tissue region. Higher LogFC values indicate better tissue specificity and decontamination accuracy. **c**, Spatial distribution of transcript counts. The top and bottom rows display the spatial expression patterns of mouse and human transcript counts, respectively, across the Original, SpotClean, and DiffGSP. d, ROC curves evaluating the performance of using species-specific transcript counts to classify spot identity, with Area Under the Curve (AUC) values reported for each method. **e**,**f**, Performance comparison on simulated datasets across 15 samples evaluated by (**e**) Pearson Correlation Coefficient (PCC) and (**f**) Mean Squared Error (MSE). Each boxplot illustrates the distribution of metrics calculated across all simulated genes. **g**, Spatial expression maps of four representative genes (Gene-1 to Gene-4) demonstrating the capacity of each method to recover the ground-truth, pre-diffusion spatial patterns.

We further evaluated species classification performance in section S2 by leveraging human- and mouse-specific transcript abundances to assign species labels to individual spots and quantified classification performance using the area under the receiver operating characteristic curve (AUC) (**Fig. 2d**). DiffGSP achieved improved species separation, further demonstrating its ability to reduce diffusion-induced transcript mixing. To determine whether conventional spatial denoising approaches could also correct molecular diffusion, we applied Sprod and STAGATE^40^ to the same datasets. Unlike diffusion-aware approaches such as DiffGSP and SpotClean, both methods introduced pronounced smoothing effects that blurred spatial patterns (**Supplementary Fig. 5a–c**). In some regions, these methods even reduced spatial specificity compared with the original data, resulting in lower LogFC values (**Supplementary Fig. 5d**). These results demonstrate that molecular diffusion correction is fundamentally distinct from conventional spatial denoising and requires dedicated modeling of the underlying diffusion process.

Next, we evaluated DiffGSP using simulated datasets with known ground truth, as the true pre-diffusion expression patterns cannot be directly observed in real spatial transcriptomics data. We generated spatial expression landscapes using simstpy^41^ and introduced physically modeled molecular diffusion together with technical noise to mimic real ST data. To assess robustness, datasets with varying tissue sizes and multiple replicates were generated (see **Methods** for details). Compared with the original diffusion-distorted data, both DiffGSP and SpotClean significantly improved recovery accuracy, as indicated by increased Pearson correlation coefficient (PCC) and reduced mean squared error (MSE) relative to ground truth (**Fig. 2e–g**). Among all evaluated methods, DiffGSP achieved the best overall performance, demonstrating its superior ability to reconstruct diffusion-distorted spatial expression profiles. In contrast, Sprod and STAGATE failed to recover diffusion-induced distortions (**Supplementary Fig. 6a–c**).

Beyond spatial localization, molecular diffusion can also distort gene–gene relationships and potentially bias downstream network analyses (**Extended Data Fig. 1a**). We therefore examined whether DiffGSP could preserve gene correlation structures after diffusion correction. Comparison of PCC values between gene pairs across different data sources showed that DiffGSP more faithfully recovered the ground-truth correlation structure (**Extended Data Fig. 1b**). For representative gene pairs, both the original data and SpotClean exaggerated positive and negative correlations (**Extended Data Fig. 1c–d**), potentially introducing spurious associations in downstream analyses. In contrast, DiffGSP effectively reduced this distortion and preserved correlation patterns consistent with the ground truth, improving the reliability of downstream biological interpretation.

Since molecular diffusion can generate smooth gradient-like patterns, we further investigated whether inverse diffusion reconstruction would distort naturally occurring spatial gradients. Four representative gradient expression patterns were simulated, followed by diffusion simulation and recovery using different methods (**Extended Data Fig. 2a**). DiffGSP accurately reconstructed the original gradients while preserving their intrinsic spatial structures (**Extended Data Fig. 2a–b**), indicating that it does not introduce artificial sharpening or distort biologically meaningful spatial trends. Given the numerical sensitivity of inverse diffusion, we further evaluated the stability of diffusion coefficient selection. Across a range of candidate diffusion coefficients, the stability indicator exhibited a clear transition from effective correction to unstable amplification (**Extended Data Fig. 2c–d**). The coefficient automatically selected by DiffGSP corresponded to this transition point and achieved accurate expression recovery (**Supplementary Fig. 7a**), supporting the effectiveness of the adaptive parameter adjustment strategy in mitigating the numerical instability associated with the ill-posed inverse diffusion problem.

We further evaluated the computational scalability and parameter robustness of DiffGSP. On simulated datasets, we assessed computational efficiency in terms of runtime, memory usage, and GPU memory consumption. For small-scale datasets, DiffGSP required less computational time and memory than SpotClean (**Supplementary Fig. 8a–c**). For large-scale datasets, DiffGSP maintained high efficiency through its subgraph-based reconstruction strategy (**Supplementary Fig. 8d–f**; see **Methods**). We further performed sensitivity analyses on three key parameters: the initial diffusion coefficient, which controls the strength of diffusion correction and is subsequently adjusted by DiffGSP’s adaptive strategy to mitigate numerical instability during inverse reconstruction; the low-pass filtering coefficient, which stabilizes inverse reconstruction by suppressing noise; and the iteration step, which determines the effective backward diffusion time. DiffGSP showed robust performance across a wide range of parameter settings, demonstrating the stability of its reconstruction framework (**Supplementary Fig. 9a–b**).

Overall, these results demonstrate that DiffGSP accurately and robustly reconstructs diffusion-distorted spatial expression profiles while maintaining computational scalability and parameter robustness. In the following sections, we further investigate how the restored spatial fidelity improves downstream biological analyses.

### DiffGSP restores spatial gene expression fidelity and enables accurate tissue architecture characterization across diverse spatial transcriptomics platforms

We systematically evaluated DiffGSP across multiple sequencing-based spatial transcriptomics platforms in mouse brain, including ST, 10x Visium, Slide-seqV2, and Stereo-seq (**Fig. 3a**). The mouse brain provides an ideal system for evaluating spatial fidelity because its highly organized anatomical structures require accurate preservation of spatial boundaries and molecular distributions. Anatomical regions were annotated based on the Allen Mouse Brain Atlas^42^ (**Fig. 3b**).

**Fig. 3.**
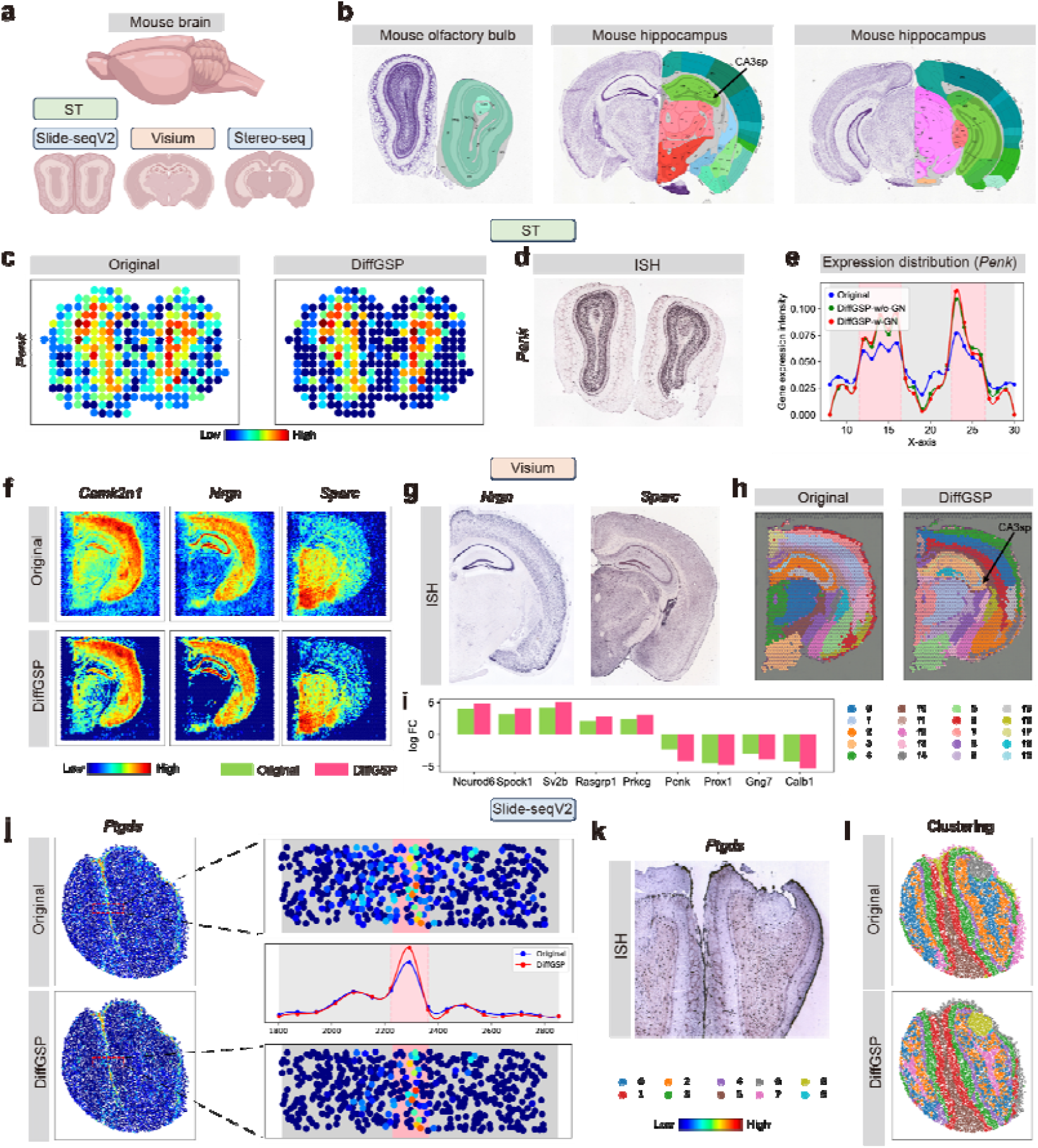
DiffGSP restores spatial expression patterns and improves tissue architecture characterization across diverse mouse brain spatial transcriptomics datasets. **a**, Schematic overview of the mouse brain spatial transcriptomics datasets used for validation, including ST, Visium, Slide-seqV2, and Stereo-seq. **b**, Reference anatomical annotations of the mouse olfactory bulb and hippocampus from the Allen Mouse Brain Atlas. **c**, Comparison of spatial expression patterns for the *Penk* expression in the ST dataset between the original and DiffGSP-processed data. **d**, ISH reference image from the Allen Mouse Brain Atlas validating the spatial localization of *Penk* expression. **e**, Spatial gene expression intensity profile of *Penk* sampled along the horizontal coordinate (x-axis), comparing the original data with DiffGSP variants with or without gene network smoothing (-w/o-GN and -w-GN). **f**, Spatial expression distributions of three representative genes (*Camk2n1, Nrgn*, and *Sparc*) in the 10x Genomics Visium dataset. **g**, Corresponding ISH reference images for *Nrgn* and *Sparc*. **h**, Spatial domain clustering results for the 10x Genomics Visium dataset. DiffGSP processing improves the recovery of hippocampal substructures and resolves subtle anatomical regions, including the CA3sp layer (indicated by the arrow), which is blurred or misclassified in the original data. **i**, Bar plot showing the log-fold change (LogFC) of representative marker genes between identified tissue domains. DiffGSP enhances the separation of spatially adjacent regions by increasing the expression differences of region-specific marker genes. **j**, Spatial distribution of the *Ptgds* gene in the Slide-seqV2 dataset. The magnified insets demonstrate that DiffGSP restores sharper spatial expression patterns compared with the diffusion-distorted original data. **k**, Reference in situ hybridization (ISH) image from the Allen Mouse Brain Atlas illustrating the spatial localization of *Ptgds* expression. **l**, Spatial domain clustering results for the Slide-seqV2 dataset. DiffGSP improves the preservation and characterization of fine-grained spatial domains.

We first applied DiffGSP to mouse olfactory bulb (MOB) spatial transcriptomics dataset, which exhibits a well-organized laminar structure^43–45^. Using *Penk* as a representative gene and a marker of the granule cell layer (GCL) (**Fig. 3c**), we observed that the original signal shows noticeable blurring and inter-region signal leakage, whereas DiffGSP restores sharper spatial domain boundaries and more coherent spatial patterns. This improvement is supported by in situ hybridization (ISH) data (**Fig. 3d**), which shows better agreement with the expected anatomical localization. Expression intensity along a spatial axis (**Fig. 3e**) reveals a more structured and biologically plausible distribution. Consistently, clustering analysis shows improved delineation of the GCL, achieving higher agreement with CARD-defined annotations as measured by the Jaccard index^46^ (**Extended Data Fig. 3a–c**). Notably, both variants of DiffGSP (with and without gene network information) improve performance, while gene-level graph information provides additional molecular constraints for spatial reconstruction.

We next evaluated DiffGSP on a mouse brain 10x Visium dataset to assess its ability to recover anatomically organized spatial patterns. Representative marker genes, including *Camk2n1, Nrgn*, and *Sparc* (**Fig. 3f**), showed that DiffGSP restored more localized expression patterns and preserved sharper regional boundaries compared with the original data. These recovered spatial patterns exhibited improved agreement with ISH signals (**Fig. 3g**), supporting the restoration of biologically meaningful expression localization. At the tissue organization level, spatial domain analysis demonstrated that DiffGSP improved the delineation of hippocampal subregions, particularly the CA3-sp and dentate gyrus (DG), which were not clearly separated in clustering results from the original data (**Fig. 3h**). Consistently, differential expression analysis between CA3-sp and DG revealed increased separation of region-specific marker genes after DiffGSP correction, reflected by enhanced log-fold changes (**Fig. 3i**). Compared with SpotClean, DiffGSP achieved superior recovery of both spatial gene expression patterns and hippocampal anatomical structures (**Extended Data Fig. 3d–f**).

We further evaluated DiffGSP on Slide-seqV2 mouse brain data to assess its performance under higher spatial resolution. Using *Ptgds* as a representative gene (**Fig. 3j**), DiffGSP recovered more localized expression patterns and reduced spatial blurring compared with the original data. ISH comparison (**Fig. 3k**) further confirmed improved agreement with anatomical expression localization, while spatial domain analysis demonstrated better preservation of fine-grained tissue structures (**Fig. 3l**). We next applied DiffGSP to Stereo-seq mouse brain data. Using *Hpca* as an example gene (**Extended Data Fig. 3g**), DiffGSP improved spatial expression fidelity and showed better concordance with ISH signals, particularly in regions with weak or undetectable expression in the original data. At the tissue organization level, spatial domain analysis (**Extended Data Fig. 3h** and **Supplementary Fig. 10a–c**) revealed more accurate recovery of fine anatomical structures, including CA1, DG, and adjacent subregions, consistent with previous anatomical studies^5^. Together, these results demonstrate that DiffGSP effectively preserves and reconstructs fine-scale spatial organization across high-resolution spatial transcriptomics platforms.

In summary, across multiple spatial transcriptomics platforms and resolutions, DiffGSP consistently restores spatial gene expression fidelity and facilitates tissue architecture characterization, demonstrating robust and scalable performance in reconstructing biologically meaningful spatial patterns.

### DiffGSP enables refined spatial architecture delineation and functional characterization of gene modules in mouse kidney

The mammalian kidney exhibits remarkable structural complexity, reflecting its diverse and tightly regulated physiological functions, including blood filtration, blood pressure regulation, and systemic homeostasis maintenance^47–52^ (**Fig. 4a**). To evaluate whether DiffGSP can restore spatial organization in complex tissues, we applied it to kidney datasets generated using two sequencing-based spatial transcriptomics platforms with different spatial resolutions, Visium and Stereo-seq.

**Fig. 4.**
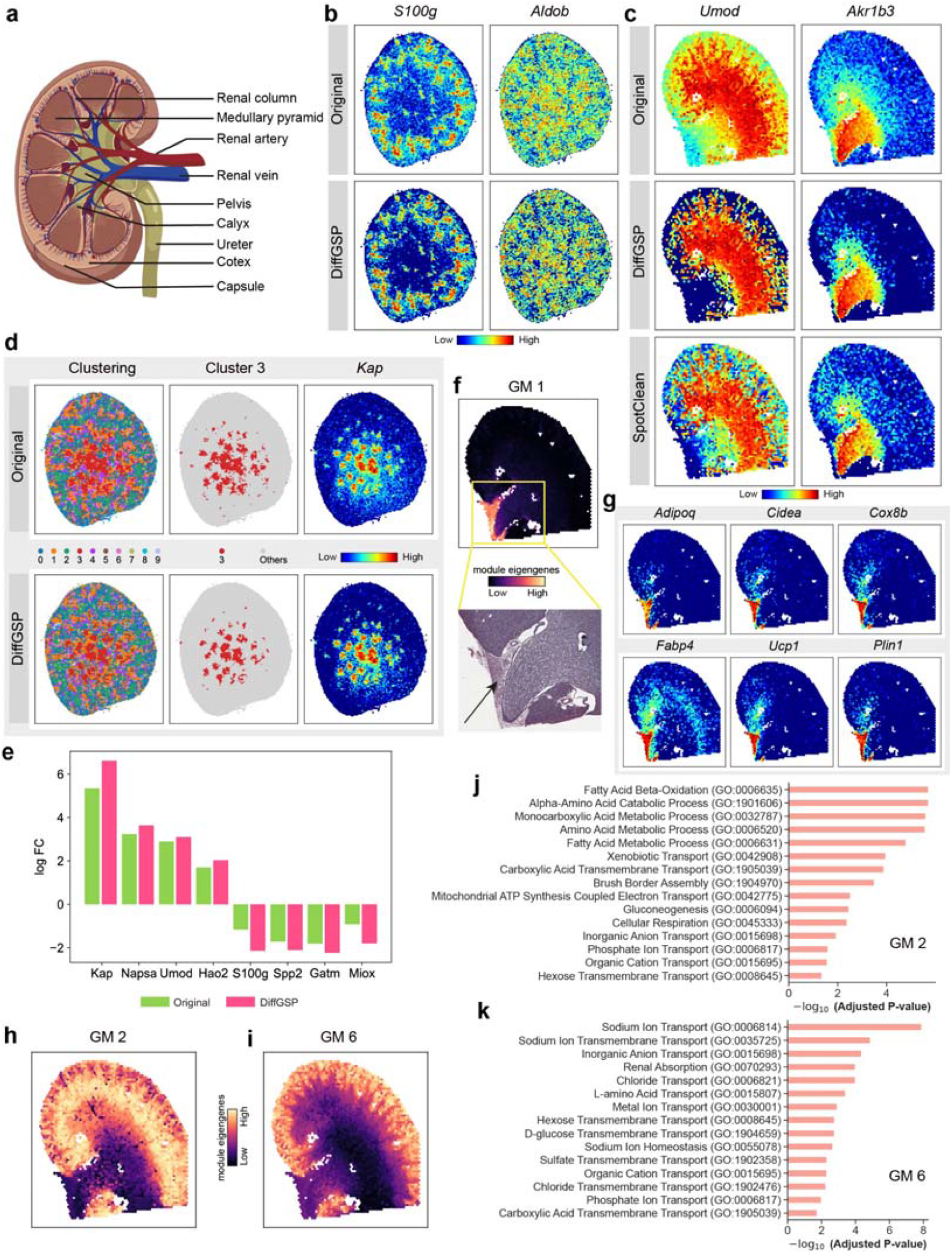
DiffGSP elucidates fine-grained anatomical structures and functional zones in the mouse kidney. **a**, Reference anatomical diagram of the mouse kidney cross-section, illustrating major structural regions including the cortex, medullary pyramid, pelvis, and capsule. **b**, Comparison of spatial expression patterns for cortex-specific genes (*S100g* and *Aldob*) between the Original and DiffGSP-processed datasets. SpotClean cannot process this dataset. **c**, Comparative spatial expression maps of medulla-associated genes (*Umod* and *Akr1b3*) across the Original, DiffGSP, and SpotClean data, highlighting the sharpened regional boundaries recovered by DiffGSP. **d**, Spatial domain clustering comparison (left panels) and isolation of Cluster 3 (middle panels) relative to its corresponding marker gene *Kap* (right panels). **e**, Bar plot showing the LogFC values of representative kidney marker genes between the Original (green) and DiffGSP-processed (pink) data. **f**, Spatial profile of Gene Module 1 (GM 1) eigengenes (top) and the aligned histological H&E staining image (bottom), with the arrow indicating the adipose region. **g**, Spatial expression patterns of six characteristic genes (*Adipoq, Cidea, Cox8b, Fabp4, Ucp1*, and *Plin1*) associated with GM 1, showing highly specific enrichment in the adipose region. **h**, Spatial distribution map of module eigengenes for Gene Module 2 (GM2), showing a broad cortical distribution extending toward the renal column. **i**, Spatial distribution map of module eigengenes for Gene Module 6 (GM6), showing preferential localization to the outer cortex. **j**, Bar plot illustrating Gene Ontology (GO) enrichment analysis results for GM 2. **k**, Bar plot illustrating GO enrichment analysis results for GM 6.

We first assessed the recovery of spatial gene expression patterns using representative kidney marker genes. In Stereo-seq data, genes such as *S100g* and *Aldob* exhibited reduced signal spreading and enhanced spatial confinement after DiffGSP processing, resulting in clearer anatomical localization compared with the original data (**Fig. 4b** and **Extended Data Fig. 4a**). Similar improvements were observed in Visium data, where genes including *Umod* and *Akr1b3* showed sharper spatial boundaries and improved regional specificity after correction (**Fig. 4c** and **Extended Data Fig. 4b**), consistent with previously reported expression patterns^53, 54^.

Spatial clustering based on DiffGSP-processed data generated more refined and anatomically coherent renal tissue domains compared with the original data (**Fig. 4d**). This improvement was supported by more specific recovery of marker gene localization. For example, *Kap*, a marker of renal proximal tubules^55^, showed a more spatially restricted expression pattern after DiffGSP correction, whereas the original data displayed broader signal distribution, likely due to diffusion-induced transcript leakage. The reduced spatial overlap after correction improved the separation of adjacent tissue domains, including cluster 1 and cluster 3 (**Fig. 4d**). Consistently, differential expression analysis revealed increased fold changes of kidney-associated marker genes between these two domains after DiffGSP correction (**Fig. 4e**), indicating enhanced regional specificity and molecular distinction.

Beyond individual genes and tissue domains, we further examined whether DiffGSP could preserve spatially organized gene co-expression structures, as molecular diffusion can distort not only spatial localization but also gene–gene relationships. We applied weighted gene co-expression network analysis using pyWGCNA^56^ to the kidney Visium dataset to identify gene modules (GMs), which consist of highly connected groups of co-expressed genes representing coordinated biological processes and spatially organized transcriptional programs. DiffGSP identified six modules, whereas only four modules were detected from both the original data and SpotClean under default settings. Moreover, module eigengenes, which summarize the overall expression patterns of genes within each module, derived from DiffGSP exhibited stronger spatial specificity and clearer correspondence with anatomical structures (**Extended Data Fig. 4c–e**). Notably, DiffGSP enabled the identification of a unique gene module (GM1) with strong spatial localization in a specific kidney region, supported by H&E staining (**Fig. 4f** and **Extended Data Fig. 4f**). This module exhibited coordinated expressions of lipid-associated genes, including *Adipoq, Cidea*, and *Fabp4* (**Fig. 4g**), suggesting a localized lipid-associated transcriptional program.

In the DiffGSP results, GM2 and GM6 were two gene modules occupying neighboring cortical regions within the kidney architecture (**Fig. 4h–i and Supplementary Fig. 11a–b**). Although both modules were primarily localized to the renal cortex and shared cortical tubular characteristics, they exhibited distinct spatial organizations: GM2 showed broader cortical distribution extending toward the renal column, whereas GM6 was preferentially localized to the outer cortex adjacent to the renal capsule. Compared with the original data and SpotClean, DiffGSP reduced spatial overlap between these modules and achieved clearer separation of fine-scale cortical structures. The spatial distribution of GM2 after DiffGSP correction was confined to the cortex, whereas the corresponding module in the original and SpotClean-processed data exhibited signal spreading into the medulla, possibly due to diffusion-induced spatial blurring. Gene Ontology (GO) analysis further supported the functional heterogeneity between GM2 and GM6 (**Fig. 4j–k**). GM2 was enriched for metabolic processes, including fatty acid and amino acid metabolism and cellular respiration (**Fig. 4j**), whereas GM6 showed stronger enrichment for ion transport and electrolyte homeostasis (**Fig. 4k**), indicating distinct functional specialization within cortical regions.

In summary, DiffGSP restores diffusion-distorted spatial gene expression patterns, enables refined delineation of kidney tissue architecture, and facilitates the identification of spatially organized gene modules. By mitigating diffusion-induced signal mixing, DiffGSP provides a more faithful representation of tissue organization and reveals functional heterogeneity that is obscured in conventional spatial transcriptomics data.

### DiffGSP more effectively clarifies tumor borders, improves cell type identification, and enhances characterization of the tumor microenvironment in human colorectal cancer

Tumors exhibit highly complex and heterogeneous spatial architectures, where diverse cellular populations dynamically interact within the tumor microenvironment. However, in sequencing-based spatial transcriptomics data, RNA molecule diffusion can blur intricate spatial patterns of genes, potentially leading to misinterpretation of tumor structure and cell-cell communication. We applied DiffGSP to multiple colorectal cancer samples across both the Visium and Visium HD platforms, to illustrate its potential to advance spatial transcriptomics studies of cancer by enabling more reliable biological discoveries.

We first applied DiffGSP to colorectal cancer Visium data and compared the gene expression patterns with both the original dataset and SpotClean-processed data. The spatial expression patterns of genes such as *CTNNB1, ID1*, and *PIGR* demonstrated that DiffGSP effectively reduced diffusion-induced spatial signal spreading, resulting in clearer tissue borders and improved contrast between adjacent regions (**Fig. 5a**). Consistent with these improvements, unsupervised clustering on DiffGSP-processed data better matched histological morphology and delineated the complex tumor architecture with clearer borders, whereas the original data showed coarser assignments, suggestive of diffusion and noise (**Fig. 5b**). Next, three histology-guided regions of interest (ROIs) were selected for detailed analysis, and tumor regions were identified based on gene expression differences of the clusters (**Fig. 5c** and **Supplementary Fig. 12a**). In ROI 1, only DiffGSP successfully revealed finer tumor substructures. In ROIs 2 and 3, both DiffGSP and SpotClean uncovered these finer structures. In contrast, the original data failed to resolve these subregions due to blurred borders caused by molecular diffusion, highlighting the critical role of mitigating diffusion effects to accurately capture fine tumor architecture. ROI 1 contained two tumor subregions, Tumor I and Tumor II. The DEGs between the two tumor subregions exhibited higher fold-change values in DiffGSP-processed data (**Fig. 5d**). Differential expression across Leiden clusters supported distinct transcriptional programs among major tumor-associated domains (**Supplementary Fig. 12b**). Furthermore, pathway analysis based on the DEGs highlighted divergent hallmark activities between the two tumor subregions, with Tumor I enriched in proliferative/metabolic pathways and Tumor II enriched in Epithelial Mesenchymal Transition pathway (**Supplementary Fig. 12c**). These findings demonstrate that DiffGSP enables more accurate delineation of tumor subregions and their underlying biological characteristics.

**Fig. 5.**
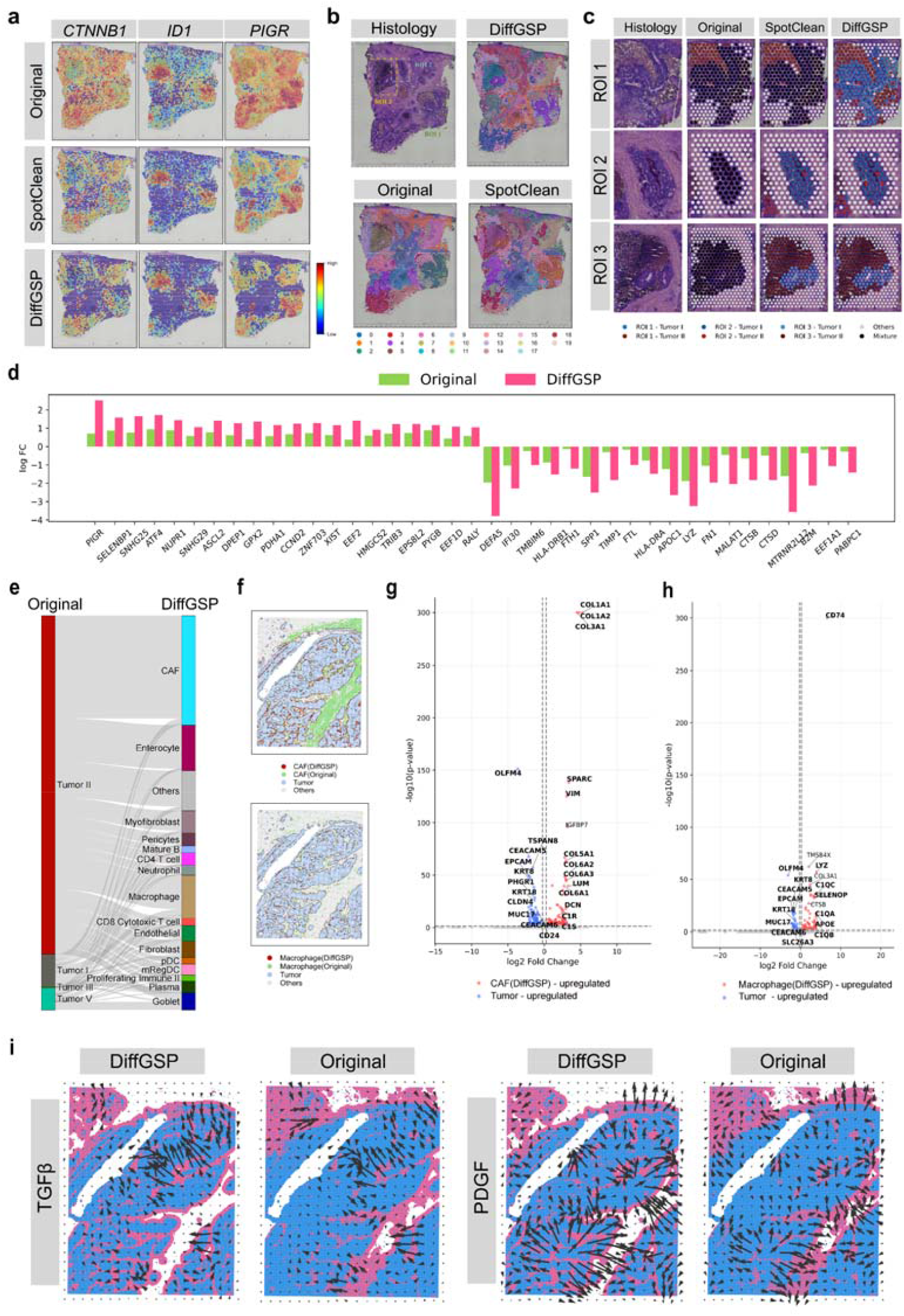
DiffGSP enhances spatial resolution of tumor architecture and refines characterization of the tumor border microenvironment in colorectal cancer data. **a**, Comparison of expression patterns of tumor marker genes in human colorectal cancer Visium data. **b**, Histology and clustering comparison among original, SpotClean-processed and DiffGSP-processed data. **c**, Tumor subtypes and border of ROI 1, ROI 2 and ROI 3 corresponding to panel **b. d**, Log fold-change values of differentially expressed genes (DEGs) between Tumor I and Tumor II of ROI 1. **e**, Sankey diagram illustrating the reassignment of cell-type annotations for regions originally labeled as tumor but reclassified as non-malignant lineages after DiffGSP processing. **f**, Spatial maps illustrating reannotation of CAFs (top) and macrophages (bottom). **g**, Volcano plot of DEGs comparing spots reannotated as CAFs by DiffGSP but labeled as tumor in the original data with tumor spots within 10 μm of the tumor-nontumor interface. **h**, Volcano plot of DEGs comparing spots reannotated as macrophages by DiffGSP but labeled as tumor in the original data with tumor spots within 10 μm of the tumor-nontumor interface. **i**, Activities of the TGF-β and PDGF signaling pathways and the inferred intercellular communication between tumor regions and tumor borders.

We then applied DiffGSP to human colorectal cancer Visium HD data^57^, considering two different spatial resolutions (8□μm and 16□μm). The cell annotations were obtained by calculating Pearson correlation coefficients with a matched scRNA-seq reference^57^. Across both resolutions, DiffGSP produced more spatially coherent cell type annotation maps that showed better concordance with histological images, particularly within tumor-enriched and Goblet-cell-enriched regions highlighted by ROIs (**Extended Data Fig. 5a, Supplementary Fig. 13a** and **Extended Data Fig. 6a**), and sharper tumor boundary maps (**Extended Data Fig. 5b, Supplementary Fig. 13b** and **Extended Data Fig. 6b**). Consistently, tumor-marker expression was elevated in tumor regions but decreased in boundary regions after DiffGSP processing (**Supplementary Fig. 13c** and **Extended Data Fig. 6c**).

To determine whether diffusion correction improves the interpretation of stromal and immune niches at the tumor-microenvironment interface, we focused on regions that were annotated as tumor in the original data but reassigned to non-malignant lineages, most prominently CAFs and macrophages (**Fig. 5e**). Spatial mapping further showed that these reassigned CAF and macrophage regions formed plausible peritumoral patterns that were less apparent in the original annotations (**Fig. 5f**). To ensure that these reassigned regions capture genuine stromal and immune programs, rather than residual tumor-derived transcriptional signal, we performed differential expression analyses using DiffGSP-derived labels, comparing (i) spots annotated as CAF (or macrophage) after DiffGSP but annotated as tumor in the original data, against (ii) tumor spots located within 10 μm radius of the tumor-nontumor interface (see “**Methods**”). CAF-reassigned regions markedly upregulated canonical extracellular matrix (ECM) remodeling genes (e.g. *COL1A1, COL1A2, COL3A1*) and other stromal markers such as *SPARC* and *VIM*. In contrast, adjacent tumor neighborhoods preferentially expressed epithelial/tumor markers including *EPCAM, CEACAM5*, and *KRT8* (**Fig. 5g** and **Extended Data Fig. 6d**). Similarly, macrophage-reassigned regions were enriched for antigen presentation and myeloid programs, exemplified by *CD74, LYZ* and complement components (*C1QA*/*C1QB*/*C1QC*) (**Fig. 5h** and **Extended Data Fig. 6d**). Consistent with these compartment-specific signals, spatial feature maps of *COL1A1* and *COL1A2* delineated CAF-enriched niches, while *CD74* highlighted macrophage-enriched territories, collectively corroborating the compartment-specific reassignments (**Extended Data Fig. 5c** and **Extended Data Fig. 6e**).

Furthermore, we examined CAF subtype signatures as a function of distance to the tumor margin. The myofibroblastic CAF (myCAF) signature peaked at the tumor-stroma interface and declined sharply with increasing distance, consistent with the preferential activation of contractile, TGF-β-associated fibroblast programs at the invasive margin. The inflammatory CAFs (iCAF) signature, by contrast, was the lowest at the boundary and progressively increased toward intermediate stromal distances, suggesting that cytokine-associated fibroblast programs predominate in regions distal from the tumor. The matrix CAF (matCAF) signature remained broadly elevated but showed a gradual decline with distance (**Extended Data Fig. 5d)**. Together, these spatial gradients support a model in which CRC stroma is spatially organized, with TGF-β and ECM-remodelling programs concentrated near the invasive margin and inflammatory fibroblast programs more prevalent in distal regions. This organization aligns with the prior reports demonstrating that CAF subtypes exhibit distinct, spatially conserved distribution patterns^58^. Finally, we assessed how diffusion correction impacts the inference of cell-cell communication. Using COMMOT, DiffGSP-processed data exhibited stronger and spatially clearer intercellular signaling across multiple essential signaling pathways, including TGF-β and PDGF (**Fig. 5i**), as well as FGF and WNT (**Extended Data Fig. 5e**).

In summary, DiffGSP improves the spatial resolution and biological interpretability of colorectal cancer transcriptomic data. By correcting diffusion-induced expression spillover at tumor borders, DiffGSP enables more precise delineation of the tumor-stroma-immune interface, refines cell type and spatial domain annotations and facilitates the recovery of compartment-specific transcriptional programs underlying stromal remodeling, immune infiltration, and intercellular communication that would otherwise be masked by diffusion-induced spatial distortion.

### DiffGSP deciphers tumor heterogeneity, tumor-stroma border and immune niche in lung cancer

Finally, we applied DiffGSP to high-resolution spatial transcriptomics data obtained through the Stereo-seq profiling of human lung adenocarcinoma (LUAD) tissue sections. Cell type annotation was performed using marker genes provided in Supplementary Table 1. Tumor architectural features, including spatial organization and cellular composition within the heterogeneous tumor microenvironment, play pivotal roles in LUAD diagnosis and clinical outcomes. After applying DiffGSP to the data, transcriptomic signal contamination attributable to spatial diffusion artifacts was markedly attenuated in tumor-adjacent non-tumor regions (**Supplementary Fig. 14a, Extended Data Fig. 7a**, and **Extended Data Fig. 8a**). Comparative analysis demonstrated that DiffGSP significantly enhanced the delineation of tumor borders compared to the original data (**Fig. 6a** and **Extended Data Fig. 7b–c**, and **Extended Data Fig. 8b–c**), thereby enabling more precise characterization of tumor–stroma interactions within the tumor microenvironment. Differential expression analysis between the tumor boundary and the tumor core revealed a substantially richer and more boundary-specific transcriptional program after DiffGSP processing, with limited overlap with the DEG set obtained from the original data. Importantly, the genes uniquely highlighted by DiffGSP recovered biologically coherent gene signatures characteristic of the tumor-stroma interface, including prominent immune-associated transcriptional programs (**Extended Data Fig. 7d** and **Extended Data Fig. 8d**).

**Fig. 6.**
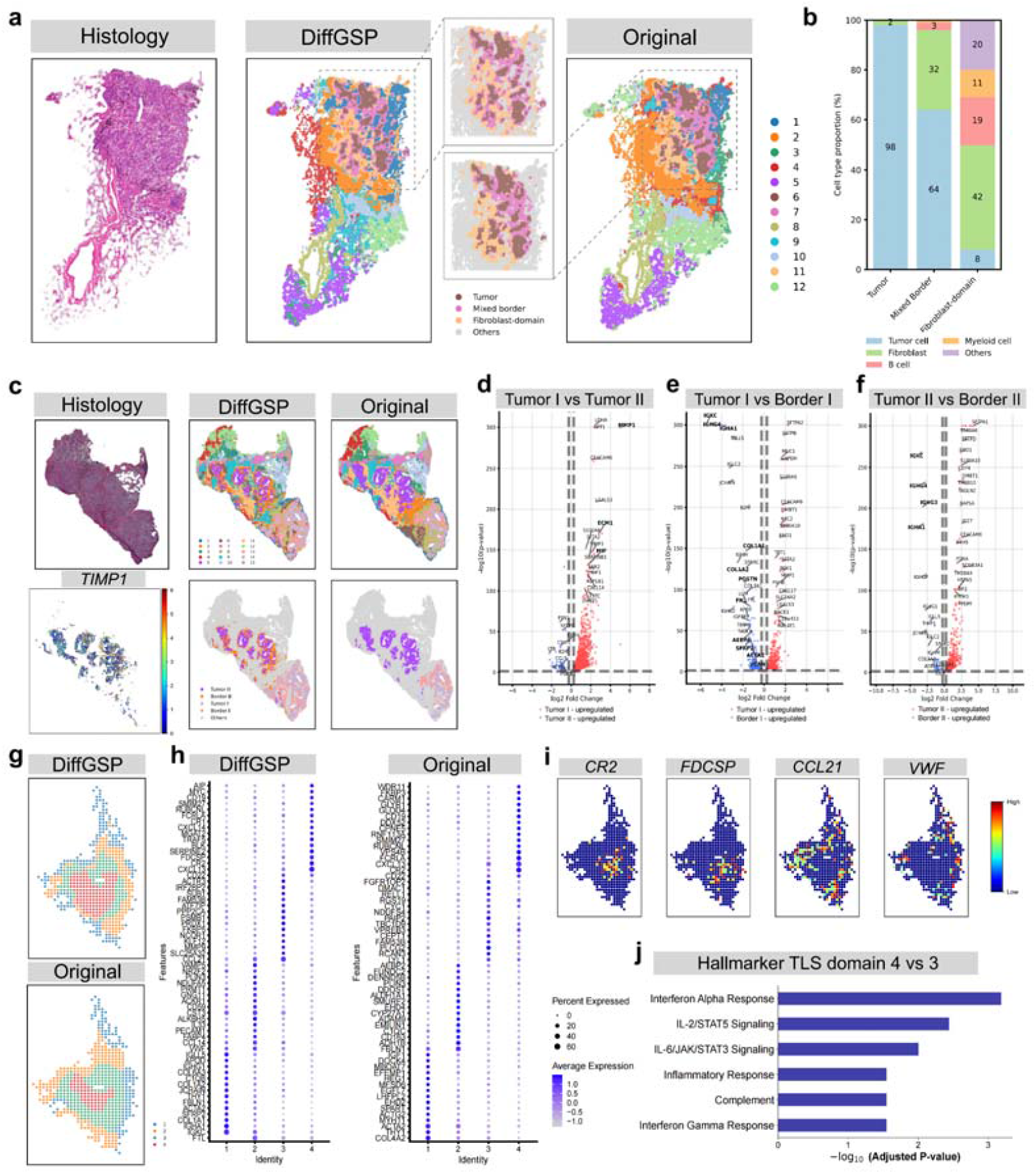
DiffGSP deciphers tumor heterogeneity, tumor-stroma border and immune niche in lung cancer. **a**, Histology and clustering comparison between original and DiffGSP-processed data in sample P1 of human lung cancer Stereo-seq data. The tumor-stroma border structures are highlighted. **b**, Cell type proportion of tumor, mixed border, and fibroblast-domain for DiffGSP-processed data. **c**, Histology and clustering comparison between DiffGSP-processed and original data, and marker gene TIMP1 of Border II in tissue sample P4. The tumor spatial heterogeneity structures are highlighted. **d**, Volcano plot showing DEGs between Tumor I and Tumor II from the DiffGSP-processed dataset. **e**, Volcano plot showing DEGs between Tumor I and Border I from the DiffGSP-processed dataset. **f**, Volcano plot showing DEGs between Tumor II and Border II from the DiffGSP-processed dataset. **g**, Clustering comparison of tertiary lymphoid structure (TLS) between DiffGSP-processed and original data in sample P4. **h**, Dotplot showing the average expression and expressed percentage of DEGs identified in TLS across all clusters in the DiffGSP-processed data and original data. **i**, Gene expression patterns of marker genes in mature TLS. **j**, Hallmark activities of DEGs between domain 4 and domain 3 corresponding to **g**.

We further performed spatial domain detection and assessed cell type distribution gradients across key anatomical regions, including the tumor core, invasive margin, and peripheral stromal compartments. Spatial profiling uncovered a distinct pattern of tumor cellular organization. The fibroblasts and myeloid lineage cells exhibited a gradual increase in abundance with increasing distance from the tumor epicenter, indicating a peripheral enrichment gradient. In contrast, malignant cells were predominantly enriched within the tumor core, consistent with a centrally confined distribution pattern (**Fig. 6b**). Intriguingly, in the original dataset, the fibroblast domain retained an appreciable tumor fraction, consistent with diffusion-driven spillover of malignant transcripts into adjacent non-tumor regions (**Supplementary Fig. 14b**), likely due to signal diffusion artifacts.

Furthermore, leveraging DiffGSP processing, we identified two distinct tumor subtypes, each associated with a specific tumor niche (**Fig. 6c**). The two tumor subtypes displayed distinct molecular profiles indicative of differential malignant potential. Tumor I exhibited strong upregulation of pro-metastatic effectors, including MMP1, MIF, and ECM1, suggestive of a more aggressive and invasive phenotype compared to Tumor II (**Fig. 6d**). These phenotypic differences may partly reflect divergent stromal and immune cell compositions at the respective tumor-stroma interfaces. Moreover, Border I demonstrated significantly upregulated expressions of extracellular matrix remodeling, cell adhesion and migration-related genes, *COL1A1, COL1A2, FN1* and *POSTN*, as well as pro-metastatic genes, *VCAN, SFRP2, ACTA2*, and *AEBP1* (**Fig. 6e**). Notably, a region termed “Border II”, which was undetectable in the original data, emerged in the DiffGSP-refined analysis. This region exhibited marked overexpression of *IGHA1, IGHG3, IGHG4* and *IGKC*—immunoglobulin heavy and light chain genes reflecting active B cell differentiation and antibody production, suggesting an active role of B cell-mediated adaptive immunity in tumor cell elimination (**Fig. 6f**). This further indicated an increased infiltration of immune cells in the Border II area surrounding Tumor II, potentially contributing to antitumor immune surveillance and malignant cell elimination. Similarly, comparative analysis of Border I and Border II revealed distinct gene expression patterns, further corroborating our previous findings **(Supplementary Fig. 14c–d**).

DiffGSP processing also uncovered distinct spatial characteristics of tertiary lymphoid structures (TLSs), ectopic lymphoid aggregates that arise within non-lymphoid tissues and comprise organized clusters of B cells, T cells, and dendritic cells. The functional competence of TLS is closely linked to their maturation state. Maturation is governed by both spatial architecture and immune cell composition, which collectively determine their capacity to coordinate antitumor immune responses. In the LUAD section, DiffGSP significantly enhanced the spatial resolution and structural clarity of mature TLS (**Fig. 6g**). The clustering results derived from DiffGSP processing exhibited sharper spatial organization, accompanied by markedly elevated expression of canonical marker genes (**Fig. 6h**). Spatial mapping revealed a compartmentalized TLS architecture: *CR2* and *FDCSP* were co-localized in the core, marking a mature follicular dendritic cell (FDC) network that supports antigen retention and B cell selection. In contrast, *CCL21* was enriched around the core, produced predominantly by fibroblastic reticular cells that orchestrate T cells and dendritic cells during TLS maturation. *VWF* expression was mainly peripheral, indicating adjacent vascular structures that may facilitate circulating lymphocyte entry (**Fig. 6i**). Hallmark enrichment analysis revealed significant enrichment of IL-2/STAT5 signaling, IL-6/JAK/STAT3 signaling, the complement system, and interferon-response programs within the TLS core. While these programs collectively delineate an immunologically active niche, they reveal a spatially orchestrated balance: IL-2/STAT5 signaling primarily sustains T-cell fitness and expansion, whereas the interferon programs drive robust antigen presentation and Th1-polarized immune activation. IL-6/JAK/STAT3 signaling indicates a cytokine-responsive microenvironment that may promote lymphocyte survival, B-cell activation, and local inflammatory maintenance. Notably, the enrichment of complement components aligns perfectly with the core-localized CR2^+^/FDCSP^+^ FDC network, which functions in antigen retention and promotes germinal-center B-cell selection. This unique functional and architectural profile strongly supports the definition of cluster 4 as a mature, immunologically active germinal center-like compartment (**Fig. 6j**).

In summary, DiffGSP not only provides a clear analysis of tumor heterogeneity, tumor-stroma border, and immune niche characteristics in the tumor microenvironment of lung cancer, but also reveals the distribution patterns, subtype molecular features, and functional status of tertiary lymphoid structures within the tumor microenvironment. These findings provide new perspectives and a valuable framework for deeper mechanistic understanding of the tumor immune microenvironment and for informing immunotherapy strategies in lung cancer.

## Discussion

Spatial transcriptomics has transformed the study of tissue organization by enabling simultaneous profiling of gene expression and spatial localization. The spatial arrangement of diverse cell types and molecular programs provides essential insights into tissue function and biological system behavior. However, sequencing-based spatial transcriptomics platforms remain vulnerable to molecular diffusion during transcript capture, which introduces spatial signal mixing and compromises the fidelity of measured expression patterns. Importantly, molecular diffusion represents a physical distortion process rather than a conventional form of technical noise, and therefore requires explicit modeling and correction to recover the underlying spatial gene expression landscape.

In this study, we developed DiffGSP, a physics-informed framework that addresses molecular diffusion in spatial transcriptomics through the integration of Fick’s law with graph signal processing. Unlike existing approaches that primarily rely on statistical assumptions, smoothing strategies, or data-driven heuristics, DiffGSP explicitly models the biophysical diffusion process and reformulates spatial transcriptomic recovery as a graph-based inverse diffusion problem. Importantly, DiffGSP distinguishes diffusion correction from noise reduction as two complementary but fundamentally different tasks. The inverse diffusion solver reconstructs spatial expression signals distorted by mRNA redistribution, whereas graph spectral filtering provides additional regularization to improve reconstruction stability and suppress residual technical noise. Across diverse spatial transcriptomics platforms, spatial resolutions, and tissue types, DiffGSP consistently improved spatial fidelity and enabled more reliable biological analyses, including tissue architecture delineation, gene relationship characterization, cell-type annotation, tumor heterogeneity assessment, and cell–cell communication inference. These results demonstrate that incorporating physical principles into computational frameworks provides an effective strategy for recovering biologically meaningful signals from distorted spatial transcriptomics measurements.

Despite these advantages, several limitations remain. First, DiffGSP is currently established under an idealized assumption that mRNA redistribution during transcript capture is predominantly governed by diffusion within an approximately planar tissue section. In practice, tissue sections may exhibit uneven morphology, folding, or incomplete contact with capture arrays, and fluid movement during sample processing may introduce convective effects that are not explicitly modeled. Future extensions incorporating additional spatial priors, such as histological images, may provide tissue-aware constraints to refine spatial graph construction and improve diffusion modeling under more complex tissue environments.

Furthermore, although graph-based low-pass filtering in DiffGSP provides effective regularization to stabilize inverse diffusion and reduce technical noise, it remains a general spectral filtering strategy and may not fully capture the complex characteristics of noise and dropout patterns inherent to spatial transcriptomics data. Developing more adaptive noise-aware models or integrating advanced statistical and data-driven approaches may further enhance signal recovery while preserving biologically meaningful spatial variation.

In addition, the effectiveness of gene-level spectral filtering depends on the reliability of gene–gene relationships inferred from spatial transcriptomics data. Lower-resolution platforms generally capture broader transcriptional programs within individual spots, providing more robust gene co-expression structures and greater benefits from gene-level filtering. In contrast, high-resolution datasets with smaller capture units and increased sparsity may provide limited information for constructing accurate gene networks. Future improvements incorporating external biological knowledge or adaptive gene graph construction strategies may further enhance gene-level signal refinement across diverse spatial transcriptomics technologies.

Overall, DiffGSP establishes a physics-informed graph signal processing framework that moves beyond conventional denoising approaches by explicitly addressing molecular diffusion as a fundamental source of spatial distortion. By separating physical signal recovery from technical noise suppression, DiffGSP provides a principled strategy for improving the fidelity and interpretability of sequencing-based spatial transcriptomics data, thereby facilitating more accurate investigation of tissue organization and biological function.

## Methods

### Data preprocessing

DiffGSP accepts gene expression profiles generated from diverse spatial transcriptomics platforms as input, enabling broad applicability across sequencing-based technologies, including Visium, Slide-seq, and Stereo-seq datasets. For each dataset, the top 2,000 spatially variable genes (SVGs) ranked by SpaGFT, a graph signal processing-based method developed in our previous work, are selected for subsequent processing and parameter estimation.

### Network construction

We construct a spatial network based on the proximity between spots using their spatial coordinates. Specifically, the undirected graph *G*_*spot*_ = (*V, E*) is constructed from the spatial coordinates, where *V*= {*vi*}represents the set containing *n* spots, and *E*= {e_ij_} represents the spatial adjacency relationships between spots. An edge *e*_*ij*_ ∈ *E* is established if spot *i* and spot *j* are spatial neighbors, for example according to a distance threshold or a K-nearest-neighbor rule.

To characterize diffusion-related spatial interactions, we assign edge weights based on the reciprocal of the Euclidean distance between neighboring spots. This weighting strategy reflects the assumption that spatially closer spots exhibit stronger coupling during molecular diffusion. The corresponding adjacency matrix *A*_*spot*_ = (*w*_*ij*)_ is defined as follows:

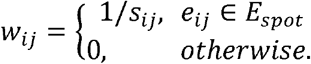

where *s*_*ij*_ is the Euclidean distance between spot *i* and spot *j*. Its corresponding degree matrix is 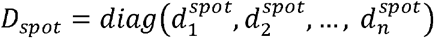, where element 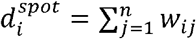 and *n* is the number of spots. The corresponding Laplacian matrix *L*_*spot*_ of graph *G*_*spot*_ is calculated from the degree matrix and the adjacency matrix as follows:

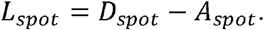

For the gene-level graph, we construct a gene co-expression network to characterize relationships among genes. The Pearson correlation coefficient is used to quantify expression similarity between gene pairs. The corresponding adjacency matrix *A*_*gene*_ = (*z*_*ij*_) is defined as follows:

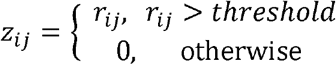

where *r*_*ij*_ is the Pearson correlation coefficient between gene *i* and gene *j*. Its degree matrix 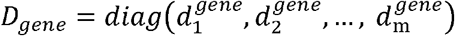, where element 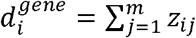 and *m* is the number of genes. The corresponding gene-level graph Laplacian *L*_*gene*_ is calculated from the degree matrix and adjacency matrix as follows:

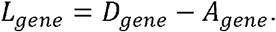

To improve numerical stability during graph signal processing, the Laplacian matrices are normalized as:

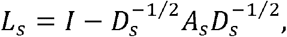

where *s* ∈ {*spot, gene*}. After normalization, all eigenvalues of *Ls* lie within the range [0, 2], facilitating stable spectral analysis^59^. The normalized Laplacians are used throughout the subsequent analysis and are denoted as *L*_*spot*_ and *L*_*gene*_ for simplicity.

### Spectral-domain filter design

#### Inverse diffusion filter

Let *f* denote the molecular concentration function of a given transcript species. Molecular diffusion is modeled according to Fick’s second law^60^:

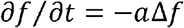

where ∂*f*⁄∂*t* describes the temporal change in molecular concentration; Δ denotes the Laplacian operator and *a* is the diffusion coefficient. Because spatial transcriptomics measurements are discrete rather than continuous, we represent the concentration field by a gene expression vector defined over spatial locations and approximate the continuous Laplacian operator using the graph Laplacian matrix *L*_*spot*_.

This discrete formulation enables the diffusion process to be modeled using graph signal processing. Let *x*_*p*_ ∈ ℝ^*n*^ and *x*_*p*+1_ ∈ ℝ^*n*^denote the gene expression vectors at diffusion steps *p* and *p*+ 1, respectively. Using a forward-difference approximation of the temporal derivative, the discrete-time form is obtained as:

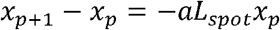

Here, *x*_*p+*1_ represents the expression state after one forward diffusion step, whereas *x*_*p*_ represents the preceding expression state. In this way, we can obtain:

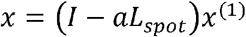

where *x*_(1)_ is the latent pre-diffusion expression vector of a gene on all spots and *x* is the observed diffused expression vector of a gene on all spots. This calculation can be further expressed in matrix form for all spots. Let *x*_(1)_,*X* E ℝ^*n*x*m*^ denote the pre-diffusion and observed gene expression matrices, respectively. In this way, we can obtain:

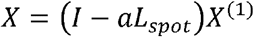

#### Low-pass filter at spot level

Let *x*_(1)_ be the output of the previous filter and *x*_(2)_ be the vector after noise removal by this filter. A Laplacian regularization term is introduced to mitigate the effects of noise. The objective function of the smoothing filter at the spot level is:

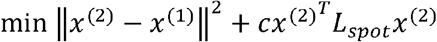

where *C* denotes the regularization coefficient of the filter. The solution to the objective function is:

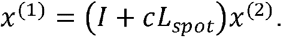

Similarly, the operation can be expressed in matrix form as:

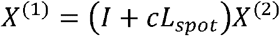

#### Low-pass filter at gene level

Let *x*^(2)^ be the output of the previous filter and *Y* be the noise-removed matrix generated by this filter. Similarly, the objective function for the smooth filter at the gene level is:

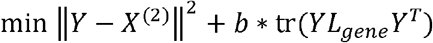

where *b* denotes the regularization coefficient of the filter and tr() denotes the trace of a matrix. The solution of the objective function is given by:

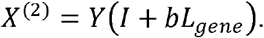

The three complementary filters are then integrated into a unified filter bank:

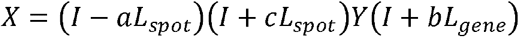

Considering multi-step diffusion and combining it with the Taubin algorithm^36^, the inverse diffusion filter and the smooth filter are iterated multiple times to obtain:

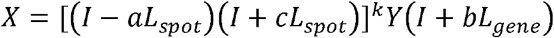

#### Filter integration

By performing spectral decomposition on the two Laplacian matrices, it is possible to avoid calculating the inverse of the matrix during iterative calculations, simplifying the calculation process. Here, we perform spectral decomposition on the Laplacian matrix ***L***_*spot*_:

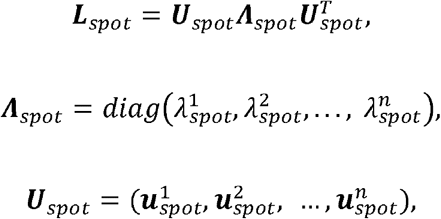

where ***Λ***_*spot*_ ∈ ℝ^*n*×*n*^ is the diagonal matrix containing the eigenvalues of ***L***_*spot*_, sorted in ascending order 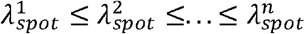, and the eigenvector matrix is ***U***_*spot*_∈ ℝ^*n*×*n*^.

Similarly, the eigenvalue matrix 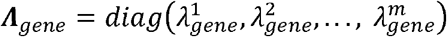 and eigenvector matrix 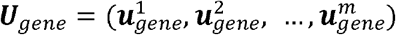 can be obtained from the graph Laplacian matrix of the gene network. For notational simplicity, ***U***_*spot*_, ***Λ***_spot_, ***U***_*gene*_ and ***Λ***_*gene*_ are denoted as ***U***_1_, ***Λ***_1_, ***U***_1_ and ***Λ***_2_, respectively.

Let *X* ∈ ℝ^n×n^ be the observed gene expression matrix and *Y* ∈ ℝ^n×n^ be the reconstructed gene expression matrix. By integrating inverse diffusion with the two low-pass filters, the final gene expression matrix is obtained as:

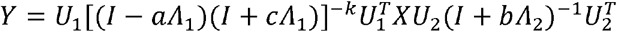

where (*I* − *aΛ*_1_)^−1^ corresponds to the inverse diffusion filter, and (*I* − *cΛ*_1_)^−1^ and (*I* − *bΛ*_1_)^−1^ correspond to the spatial low-pass filter (spot-level) and the gene-level low-pass filter, respectively.

Unlike conventional smoothing-based approaches, the inverse diffusion operator is designed to reverse the physical spreading process, whereas graph spectral filters serve as regularization operators to stabilize reconstruction and suppress technical noise.

#### Parameter estimation

We use the expression values from out-of-tissue regions in the data to estimate the three parameters: *a, b* and *c*. The objective of optimization is to make the denoised expression values outside the tissue closer to zero, and the objective function is:

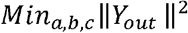

where *Y*_*out*_ represents the denoised expression matrix in out-of-tissue regions. The three parameters are constrained to be non-negative and to satisfy stability requirements of the filter, that is, 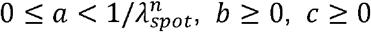. The parameters are estimated by minimizing the above objective function using the L-BFGS-B algorithm, with initial values of *a* = 0, *b* = 0 and *c* = 0. In addition, for data without background spots or that do not exhibit physical diffusion laws, we allow the three parameters to be manually adjusted.

#### Adaptive design of the graph spectral filter bank

DiffGSP adopts an adaptive filter bank configuration according to the characteristics of different spatial transcriptomics datasets. The inverse diffusion filter and spot-level spectral filter are applied to all datasets to correct diffusion distortion and suppress spatially structured noise. The gene-level spectral filter is incorporated when reliable gene–gene relationships can be inferred from the data. Specifically, low-resolution platforms with larger capture areas generally provide more robust gene co-expression patterns because individual spots capture broader transcriptional programs. Therefore, the gene-level spectral filter is applied to low-resolution datasets, including Visium. For high-resolution datasets, such as Slide-seqV2, Visium HD, and Stereo-seq, the gene-level filter is not applied due to increased sparsity and reduced reliability of gene network construction.

### Stable inverse diffusion reconstruction

During inverse diffusion, the solution becomes progressively unstable over time, reflecting the intrinsic instability of backward diffusion processes in ill-conditioned inverse problems^29, 61^. DiffGSP introduces low-pass filtering, as discussed above, to constrain the amplification of noise-sensitive high-frequency components during inverse diffusion, thereby mitigating instability in reconstruction. Another key challenge is selecting the diffusion coefficient at a fixed diffusion time, which governs the trade-off between diffusion correction and numerical stability. Mathematically, an excessively large diffusion coefficient can exacerbate the numerical instability associated with the ill-posed inverse diffusion problem, characterized by unstable amplification of reconstructed signals, where the maximum reconstructed expression value increases rapidly with increasing diffusion coefficient at a fixed diffusion time. Conversely, an excessively small diffusion coefficient leads to insufficient recovery of diffusion-distorted expression profiles.

To systematically determine an appropriate coefficient, we evaluate inverse diffusion across a range of candidate values around an initial estimate. For each diffusion coefficient setting, we compute a stability indicator defined as 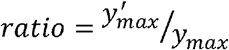, where 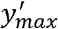 and y_*max*_ are the log-transformed maximum expression values of gene expressions after correction and before correction, respectively. This indicator can capture amplitude variations induced by inverse diffusion and an excessively large value of this indicator *ratio* typically suggests that the reconstructed solution is becoming unstable.

As the diffusion coefficient increases, this indicator exhibits a characteristic transition from a stable regime, where corrections improve gradually, to an unstable regime characterized by rapid amplitude inflation. This behavior reflects the fundamental trade-off between reconstruction fidelity and stability in inverse diffusion.

To automatically identify this transition point, we apply the Kneedle algorithm^39^ to detect the knee point of the curve. The Kneedle algorithm identifies knee points corresponding to regions of high curvature in discrete curves. The diffusion coefficient corresponding to this point is selected as the optimal parameter for DiffGSP, providing a robust balance between diffusion correction and numerical stability.

### Subgraph-based computational acceleration

To improve the scalability of DiffGSP for large-scale spatial transcriptomics datasets, we developed a spatial graph partitioning strategy for efficient inverse diffusion reconstruction. Instead of performing inverse diffusion on the entire spatial graph, DiffGSP partitions the graph into multiple local subgraphs based on spatial grids. The number and size of subgraphs are adaptively determined according to the scale of each dataset. Each subgraph is expanded with neighboring boundary regions to preserve local spatial connectivity and diffusion information. Inverse diffusion reconstruction is then performed independently within each subgraph, and the resulting profiles are integrated to obtain the final reconstructed spatial expression landscape. In this study, large-scale spatial transcriptomics datasets, including Slide-seqV2, Stereo-seq, and Visium HD datasets, were processed using this strategy. This approach substantially reduces computational and memory requirements while preserving reconstruction accuracy, enabling efficient application of DiffGSP to large-scale spatial transcriptomics datasets.

### Benchmarking on human–mouse chimeric datasets

#### Data preprocessing and data normalization

We downloaded three human–mouse chimeric spatial transcriptomics datasets containing spatial expression profiles of both human and mouse transcripts, together with spot-level annotations indicating tissue-associated regions. Transcripts were assigned to their corresponding species according to gene origin, generating separate human-transcript and mouse-transcript expression matrices for each tissue section.

For human-transcript datasets, spots annotated as human or mixed-species regions were considered tissue-associated spots, whereas all remaining spots were classified as out-of-tissue regions. Similarly, for mouse-transcript datasets, mouse and mixed-species spots were considered tissue-associated spots, and the remaining spots were treated as out-of-tissue regions.

The resulting datasets were subsequently processed using different computational methods. To ensure comparability among outputs generated by different approaches, max-value scaling normalization was applied to all processed expression matrices. Specifically, expression values were rescaled such that the maximum expression value within each dataset was set to 10,000, ensuring that datasets from different methods were placed on a comparable scale while preserving relative expression patterns.

#### Performance evaluation

To quantify transcript-level specificity across species, we calculated the log-fold change (LogFC) of average transcript abundance between species-specific regions. For human transcripts, LogFC was calculated as:

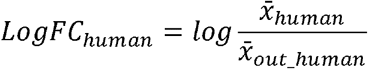

where 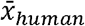 and 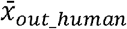 represent the average abundance of human transcripts in human tissue spots and spots outside the human tissue region, respectively. Similarly, for mouse transcripts, LogFC was calculated as:

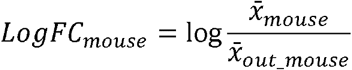

Higher LogFC values indicate stronger species-specific enrichment and improved discrimination between human- and mouse-derived transcript signals.

To further evaluate whether transcript-level specificity could improve spot-level identification, human- and mouse-derived transcript abundances were used to classify spot identities. Receiver operating characteristic (ROC) curves were generated, and the area under the curve (AUC) was calculated to quantify the discriminative capability of transcript-based classification.

#### SpotClean’s pipeline settings

To control variables, we set *gene_cutoff* = 0 in *createSlide()*, ensuring that SpotClean uses the same gene set as DiffGSP. Because SpotClean is specifically designed for Visium data, we used its default parameters for comparison. We only performed calculations on data with background spots (out-of-tissue spots). When applied to high-resolution datasets, SpotClean returned expression values for only a subset of tissue spots and required substantial computation time; therefore, its performance comparisons were limited to simulated and Visium datasets.

#### Comparison with Sprod and STAGATE

To compare DiffGSP with conventional spatial transcriptomics refinement approaches, we selected Sprod and STAGATE as representative graph-based spatial smoothing and deep learning-based reconstruction methods, respectively. Sprod was selected as a representative graph-based spatial smoothing method, whereas STAGATE was selected as a representative deep learning framework that reconstructs spatial expression patterns by leveraging graph attention mechanisms. Both methods were implemented following their official pipelines with default parameter settings. The input data and preprocessing procedures were kept consistent across methods to ensure a fair comparison.

### Benchmarking on simulation data

#### Data generation

Since the ground truth of gene expression is unknown in real datasets, we used the simstpy tool (https://github.com/pinellolab/simstpy) to generate simulated gene expressions as the ground truth, with parameters *sigma* = 10 and *n_kernels* = 20. To mimic out-of-tissue spots in real data, edge areas were added around each tissue section without gene expression. We then applied Fick’s Law to simulate diffusion on the generated expression profiles, using the finite difference method, a common technique for solving partial differential equations (PDEs) to produce data exhibiting diffusion phenomena. To better reflect real-world conditions, we introduced both noise and dropout (at a rate of 0.05). During this process, we focused on two dataset scales: 30×30 and 100×100, with different diffusion steps (5, 10, and 15 for 30×30; 10 and 20 for 100×100). Using different random seeds, we generated multiple datasets, each containing 100 spatially variable genes, to evaluate the robustness of computational methods.

#### Metrics in benchmarking

To quantify gene expression recovery accuracy in the simulated datasets, we used two metrics: PCC and MSE, where PCC measures the linear correlation and MSE measures the estimation error between the true expression values and the estimated values. For one gene,

PCC is defined as 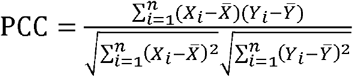 and MSE is defined as 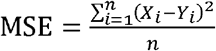, where *Y* denotes ground truth, *X* represents the DiffGSP or SpotClean estimated gene expression vector of the gene and *n* denotes the number of spots.

### Other settings in case studies

#### Differentially expressed gene analysis

The Wilcoxon rank-sum test was used to identify differentially expressed genes, with Benjamini–Hochberg-adjusted P values < 0.05 considered statistically significant. In Visium HD, we performed DEG analysis based on matched scRNA-seq to identify tumor genes.

#### Identification of spatial domains

We used the Leiden algorithm to identify spatial domains for data with neatly arranged spots. For comparisons on the same dataset, the resolution parameter was adjusted to keep the number of identified spatial domains as consistent as possible across methods. For irregularly arranged Slide-seqV2 data and Stereo-seq lung adenocarcinoma data, we used STAGATE to identify spatial domains because of their increased sparsity and spatial non-uniformity.

#### Identification of gene modules in gene co-expression network

In this analysis, we utilized the PyWGCNA package with default parameters to identify gene modules within gene co-expression networks. The *preprocess()* method was applied to prepare and normalize the data. Following this, the *findModules()* method was used to identify gene modules by grouping genes with similar expression patterns, revealing underlying biological relationships. This approach resulted in the identification of distinct gene modules in both datasets, which were further investigated for their functional significance in the context of mouse kidney analysis.

#### Cell-type identification

For Stereo-seq human lung adenocarcinoma data, cell type annotation was performed using scSorter^62^, with prior marker genes derived from matched single-cell RNA data from the same patient cohort. The proportion of each cell type was calculated within each cluster. Cell types with proportions below 0.1 within a cluster were grouped into the “others” category. Because Visium HD datasets contain a very large number of spots, PCC-based cell-type annotation was employed as a computationally efficient strategy.

#### Identification of tumor-nontumor interface and validation of reannotated cell-type regions in Visium HD data

A tumor mask was constructed by mapping tumor labels onto the spatial lattice. To reduce isolated spots and smooth local irregularities, we applied morphological operations to the two-dimensional tumor mask. Specifically, binary opening was used to remove isolated tumor regions. The tumor spots containing at least three non-tumor points in the adjacent spots were determined as the tumor-nontumor interface. To further validate that these reassigned regions capture authentic stromal and immune programs rather than residual tumor artifacts, we conducted differential expression analysis based on cell type annotations derived from DiffGSP. Specifically, we compared (i) spots reclassified as CAFs or macrophages by DiffGSP but primarily annotated as tumor in the original dataset, with (ii) tumor spots located within a 10μm radius of the tumor-nontumor interface.

#### Spatial gradient analysis of CAF subtype activities

Spots annotated as CAFs and located within 0-550 μm outside the tumor-nontumor interface were retained for analysis. Canonical signature scores for myCAF, iCAF, and matCAF were computed for each spot using *sc*.*tl*.*score_genes* (Scanpy) and subsequently standardized using Z-score. Spots were then stratified into 10 equal-width distance bins, and the mean signature score was calculated for each bin. Spatial gradients were visualized by plotting bin-wise mean scores against the distance to the tumor boundary, with cubic B-spline interpolation applied to depict continuous trends.

#### Cell-cell communication

We used COMMOT to calculate cell interactions between tumor regions and tumor borders. The analysis was performed using the default parameters specified in the official tutorial. We plotted the intercellular signaling pathways of TGF-β, PDGF, FGF, and WNT, which are closely related to tumor immunity.

#### Down-sampling, subgraph segmentation and gene set batch processing strategies

For high-resolution data, we obtained low-resolution data by merging spots. This was used for quickly and efficiently estimating parameters. A merged spot was classified as tissue-associated based on the proportion of its constituent spots located within the tissue. In order to reduce memory usage and boost speed, we partitioned the spatial graph into subgraphs based on spatial coordinates and performed DiffGSP in each subgraph, due to the fact that diffusion is fundamentally a local rather than global process. When the gene-level network was not used, genes were processed independently, allowing the gene set to be divided into batches to reduce memory usage.

## Data availability

This research was carried out using datasets that are publicly available. ST includes the MOB dataset (https://www.spatialresearch.org/resources-published-datasets/doi-10-1126science-aaf2403/)^63^. 10x Visium includes human–mouse chimeric datasets (https://www.ncbi.nlm.nih.gov/geo/query/acc.cgi?acc=GSE178221)^14^, mouse brain dataset (https://www.10xgenomics.com/datasets/mouse-brain-section-coronal-1-standard-1-1-0)^64^, mouse kidney dataset (https://www.10xgenomics.com/datasets/adult-mouse-kidney-ffpe-1-standard-1-3-0)^65^ and human colorectal cancer dataset (https://www.10xgenomics.com/datasets/human-colorectal-cancer-whole-transcriptome-analysis-1-standard-1-2-0)^66^. Slide-seqV2 includes mouse olfactory bulb dataset (https://singlecell.broadinstitute.org/single_cell/study/SCP815/highly-sensitive-spatial-transcriptomics-at-near-cellular-resolution-with-slide-seqv2#study-download)^67^. Visium HD includes human colorectal cancer datasets (https://www.10xgenomics.com/products/visium-hd-spatial-gene-expression/dataset-human-crc)^57^. Stereo-seq includes mouse brain dataset and mouse olfactory bulb dataset (https://db.cngb.org/stomics/mosta/download/)^5^, mouse kidney datasets (https://db.cngb.org/stomics/datasets/STDS0000240/data)^68^ and human lung adenocarcinoma datasets (https://ngdc.cncb.ac.cn/omix/preview/MuvMgwkc)^69^. scRNA-seq reference includes colorectal cancer dataset, matched with Visium HD (https://github.com/10XGenomics/HumanColonCancer_VisiumHD)^57^.

## Code availability

DiffGSP is implemented in Python. The source code, documentation, and usage examples will be made publicly available under an open-source license upon formal publication of the article.

## Acknowledgements

This work was supported by the National Natural Science Foundation of China (NSFC) Young Scientists Fund (Type A) (Grant No. 12625118); the Science and Technology Innovation Key R&D Program of Chongqing (Grant No. CSTB2024TIAD-STX0003); the National Natural Science Foundation of China (Grant No. 62272270); the Shandong Provincial Natural Science Foundation for Distinguished Young Scholars (Grant No. ZR2023JQ002); and the Open Project of BGI-Shenzhen, Shenzhen 518000, China (Grant No. BGIRSZ20220005). We also thank Mr. Xinyu Liu from the Second Hospital of Shandong University for his support with the analysis of human colorectal cancer data.

## Author information

These authors contributed equally: Jixin Liu, Shuli Sun and Yang Xu.

## Contributions

B.L. and X.Z. conceptualized and supervised the project. S.S. and J.L. designed and implemented the method. J.L. and S.S. implemented the simulation and benchmarking. S.S., J.L., Y.X., S.J. and S.C. collected the data. J.L., Y.X., S.S., X.Z., B.L. and G.L. analyzed the case studies and contributed to writing the manuscript. All authors reviewed and approved the final version.

## Extended Data Figures

**Extended Data Fig. 1.**
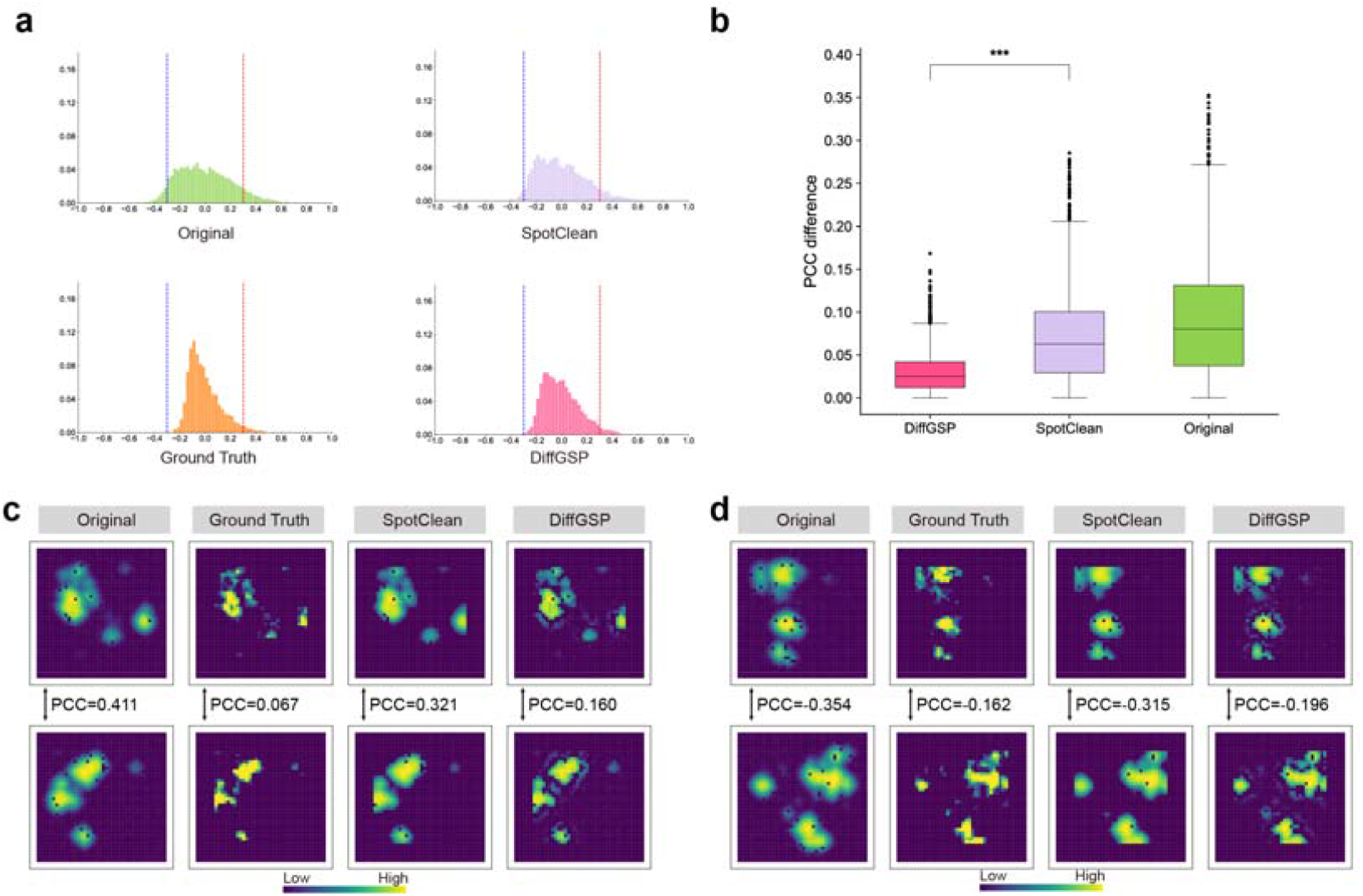
DiffGSP mitigates molecular diffusion-induced bias in gene–gene co-expression relationships. **a**, Frequency distributions of pairwise Pearson Correlation Coefficients (PCC) across all genes for the Original, Ground Truth, SpotClean, and DiffGSP datasets. Higher similarity to the Ground Truth distribution indicates a more accurate preservation of spatial gene–gene relationships. **b**, Boxplot showing the absolute PCC differences relative to the Ground Truth for each gene pair across the three methods. Significance is indicated by asterisks (p < 0.001, Wilcoxon rank-sum test). **c**, A representative gene pair demonstrates that molecular diffusion can induce false-positive spatial co-expression (high PCC), which is successfully mitigated by DiffGSP to better align with the Ground Truth. **d**, A representative gene pair demonstrates that molecular diffusion can introduce false-negative or artificially inverted spatial relationships (negative PCC), which is effectively corrected toward the Ground Truth by DiffGSP.

**Extended Data Fig. 2.**
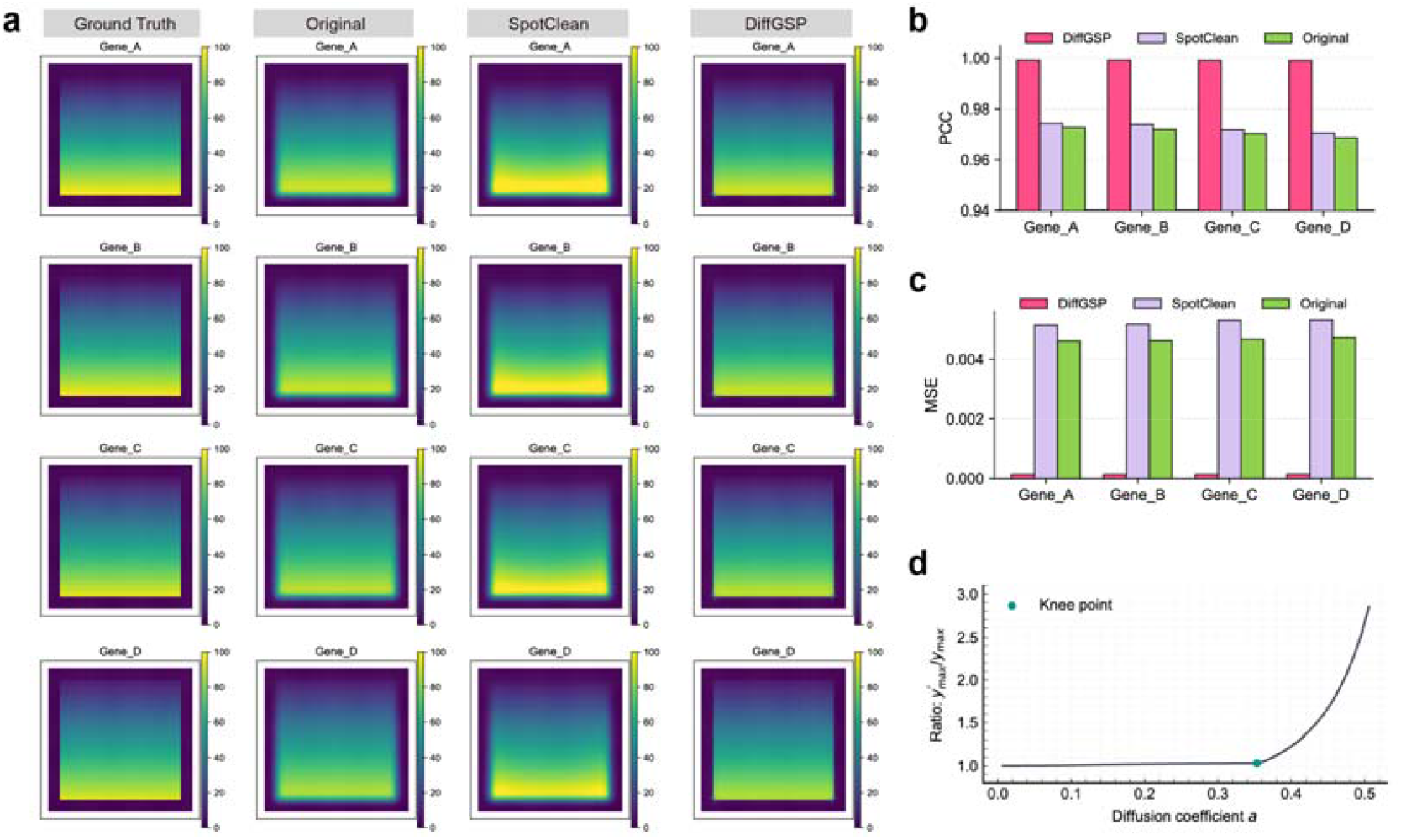
Performance of DiffGSP in recovering continuous spatial gradient patterns. **a**, Spatial expression maps of four representative genes (Gene_A to Gene_D) exhibiting continuous gradient patterns across the Ground Truth, Original, SpotClean, and DiffGSP datasets. DiffGSP accurately reconstructs the smooth spatial gradients. **b**,**c**, Performance evaluation of different methods across the four gradient genes based on **(b)** Pearson Correlation Coefficient (PCC) and **(c)** Mean Squared Error (MSE). **d**, Mathematical behavior of the decontamination process as a function of the diffusion coefficient *a*. The y-axis denotes the ratio of the log-transformed maximum expression value in the DiffGSP-processed data to that in the observed (Original) data *Ratio* = 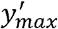/*y*_max_. The curve shows a slow, stable increase before reaching a distinct knee point (indicated by the teal dot), beyond which the ratio escalates rapidly, signifying that the reconstruction transitions into an unstable regime.

**Extended Data Fig. 3.**
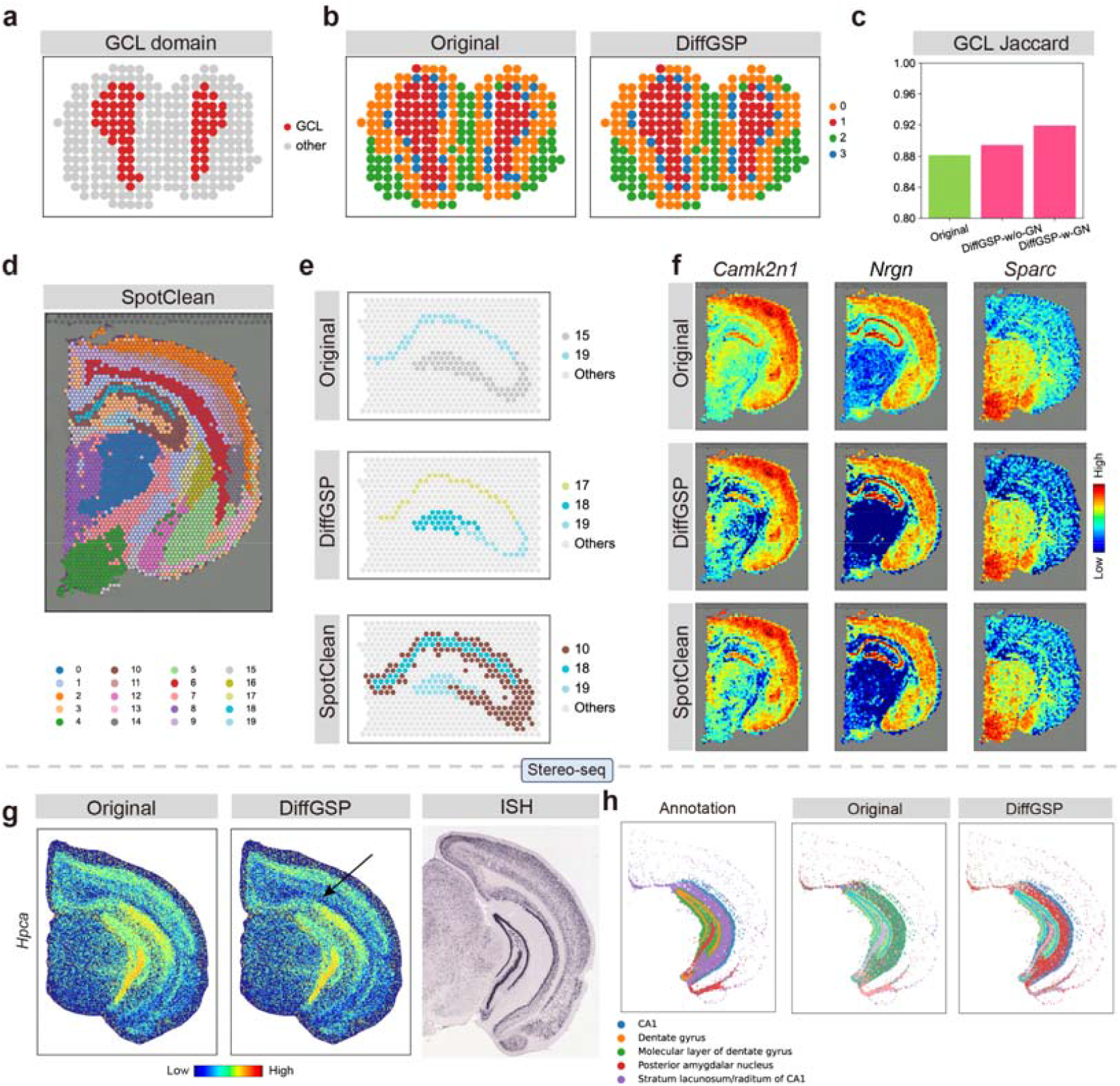
Comparative performance of DiffGSP and SpotClean in refining spatial domains and gene expression profiles. **a**, Spatial domain mask highlighting the granular cell layer (GCL) of the mouse olfactory bulb as the ground-truth reference in the ST dataset. **b**, Spatial clustering results of the ST dataset, comparing the domain definition between the Original and DiffGSP-processed data. **c**, Bar plot evaluating the clustering accuracy of the GCL domain based on the Jaccard index, comparing the original data against DiffGSP implementations with or without gene network smoothing (-w/o-GN and -w-GN). **d**, Spatial domain clustering visualization of the 10x Genomics Visium dataset processed by SpotClean. **e**, Magnified views of the reconstructed hippocampal layers (specifically clusters corresponding to the CA fields and dentate gyrus) across the Original, DiffGSP, and SpotClean datasets, highlighting the structural continuity recovered by DiffGSP. **f**, Comparative spatial expression maps of three representative genes (*Camk2n1, Nrgn*, and *Sparc*) across the Original, DiffGSP, and SpotClean Visium datasets. **g**, Spatial expression distribution of the *Hpca* gene in the high-resolution Stereo-seq dataset. DiffGSP successfully recovers a distinct, low-expression anatomical boundary layer (indicated by the arrow), matching the reference ISH characterization from the Allen Brain Atlas. **h**, Spatial clustering and domain annotations in the Stereo-seq dataset. DiffGSP aligns more faithfully with the annotation in the original study.

**Extended Data Fig. 4.**
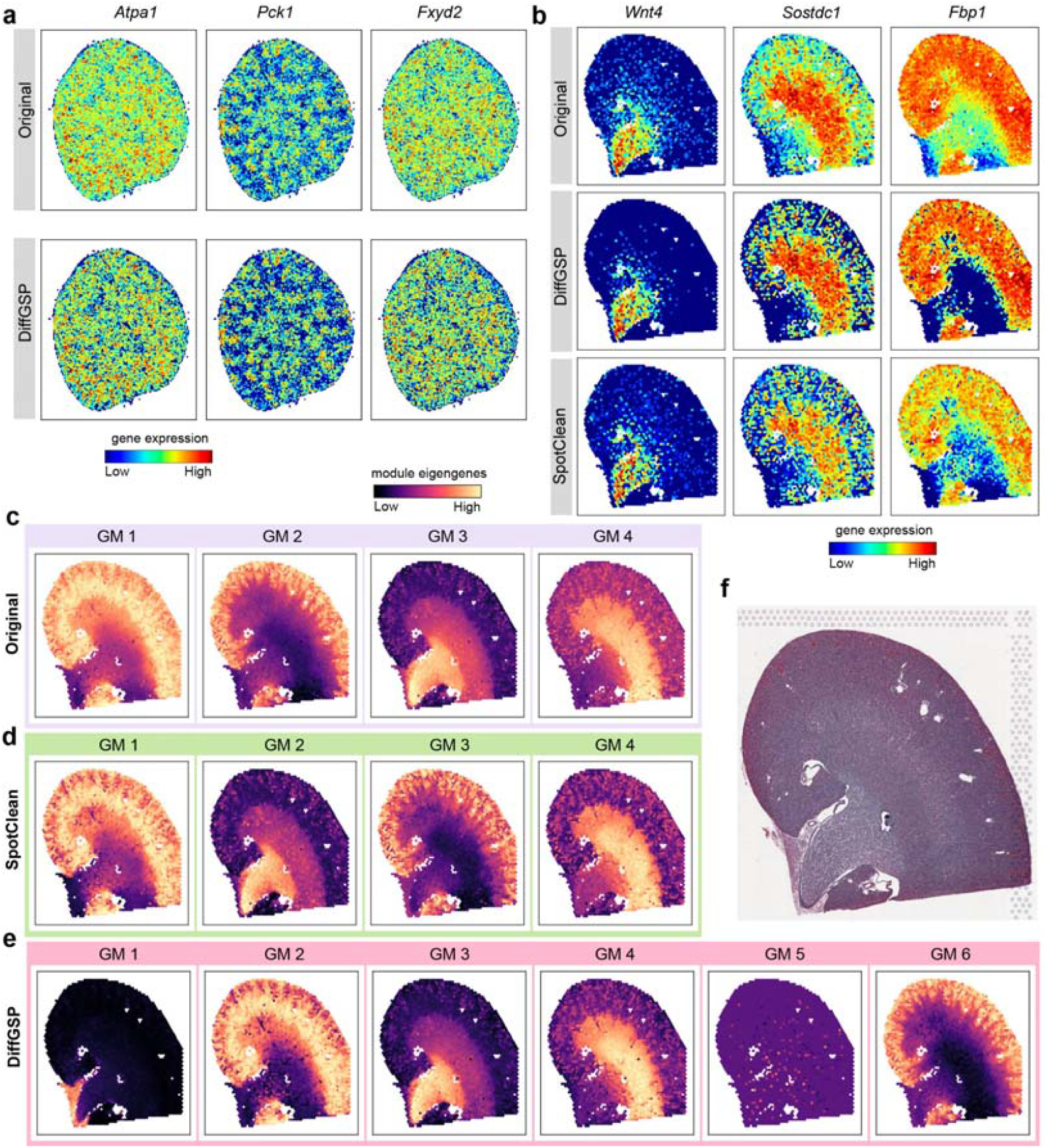
DiffGSP enhances the spatial patterns of genes and facilitates gene module identification in the mouse kidney. **a**, Comparative spatial expression maps of three representative genes (*Atpa1, Pck1*, and *Fxyd2*) in the high-resolution Stereo-seq dataset, contrasting the Original (top) and DiffGSP-processed (bottom) data. **b**, Comparative spatial expression maps of three characteristic genes (*Wnt4, Sostdc1*, and *Fbp1*) in the 10x Genomics Visium dataset across the Original, DiffGSP, and SpotClean datasets. DiffGSP demonstrates superior performance in boundary sharpening. **c**, Spatial distribution maps of module eigengenes for four Gene Modules (GM 1 to GM 4) identified in the original dataset. **d**, Spatial distribution maps of module eigengenes for GM 1 to GM 4 identified in the SpotClean-processed dataset. **e**, Spatial distribution maps of module eigengenes for the Gene Modules identified in the DiffGSP-processed dataset. Notably, DiffGSP successfully resolves two additional, spatially distinct functional modules (GM 1 and GM 5) that are obscured by molecular diffusion in the Original and SpotClean datasets. **f**, Aligned histological H&E staining image of the corresponding mouse kidney section.

**Extended Data Fig. 5.**
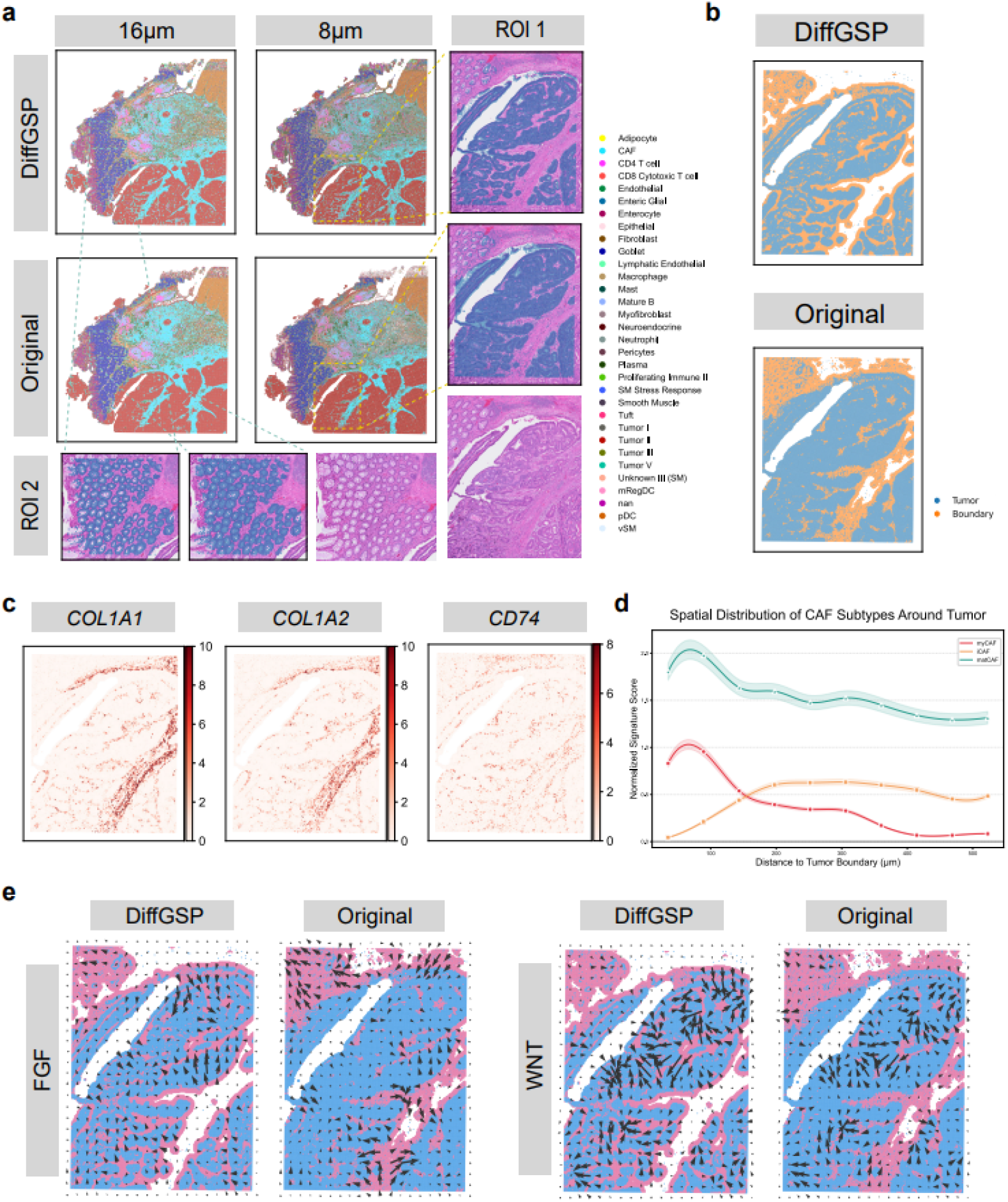
DiffGSP-enabled high-resolution spatial characterization of cell types, CAF subtype gradients, and microenvironmental interactions in colorectal cancer. **a**, Comparison of cell type identification between original and DiffGSP-processed data in human colorectal cancer Visium HD data of P1 tissue with 16μm and 8μm resolution. ROI 1 and ROI 2 represent tumor cell and goblet cell respectively. **b**, Tumor and border regions. Tumor region is defined as the region labeled with tumor cells. Tumor border is defined as the spots within 6.25 radius of the tumor region expanding outward, and some isolated spots are removed. **c**, Spatial expression patterns of CAF or macrophage representative marker genes. **d**, Spatial distribution of CAF subtypes. The x-axis indicates distance to the tumor margin. **e**, The cell-cell communication activity of FGF and WNT signaling pathways and intercellular communication between tumor region and tumor border.

**Extended Data Fig. 6.**
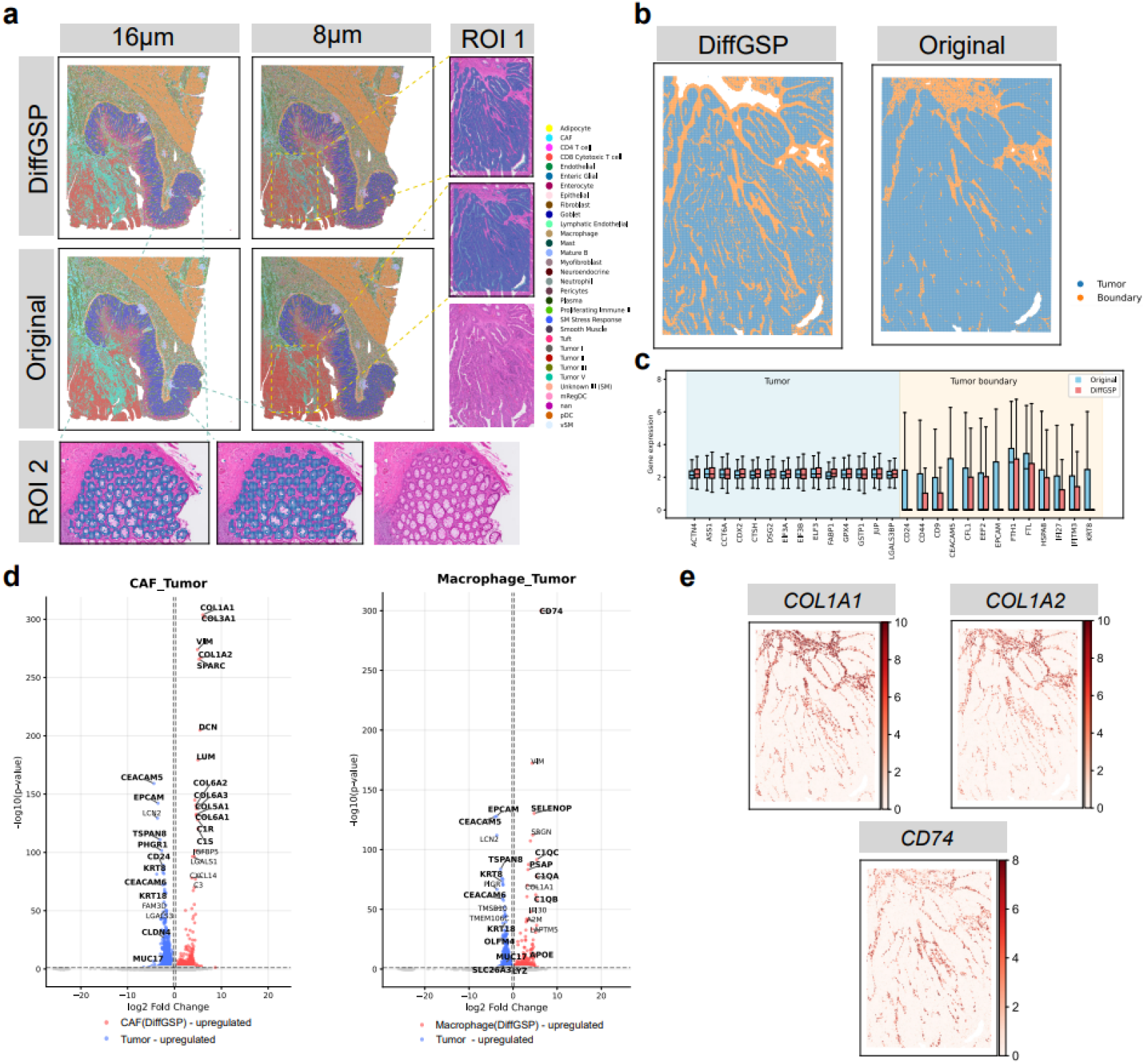
Analysis of human colorectal cancer Visium HD data of P5 tissues with 16μm and 8μm resolution. **a**, Comparison of cell-type annotations between original and DiffGSP-processed data in human colorectal cancer Visium HD data of P5 tissue with 16μm and 8μm resolution. ROI 1 and ROI 2 represent tumor cell and goblet cell respectively. **b**, Tumor and border regions. Tumor region is defined as the region labeled with tumor cells. Tumor border is defined as the spots within 6.25 radius of the tumor region expanding outward, and some isolated spots are removed. **c**, Tumor marker genes showed increased expression in the tumor region. Tumor marker genes were selected by differential expression analysis of matched scRNA-seq. Spots expressed as zero in tumor in the original data were filtered. **d**, Volcano plots of DEGs comparing spots reannotated as CAFs(left) and macrophages(right) by DiffGSP but labeled as tumor in the original data with tumor spots within 10 μm of the tumor-nontumor interface. **e**, Spatial expression patterns of CAF or macrophage representative marker genes.

**Extended Data Fig. 7.**
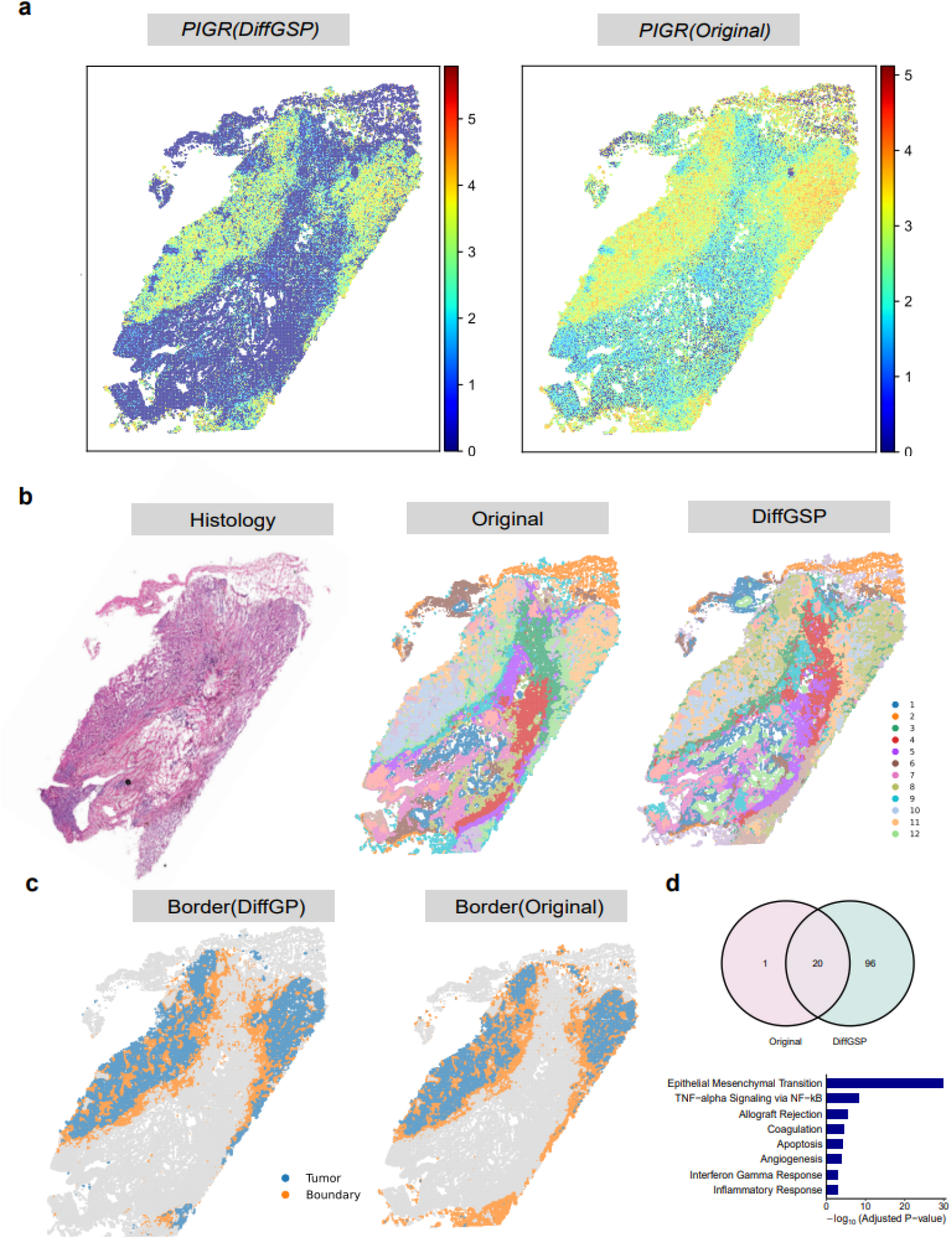
Analysis of human lung cancer Stereo-seq data of P2 tissues. **a**, Diffusion-reduced gene expression patterns in DiffGSP-processed compared to original data in sample P2. **b**, Histology and clustering comparison between DiffGSP-processed and original data. **c**, Tumor and border regions defined on the basis of the clustering results. **d**, Overlap of boundary-specific DEGs identified from the original and DiffGSP-processed data and hallmark activities of DEGs uniquely detected by DiffGSP.

**Extended Data Fig. 8.**
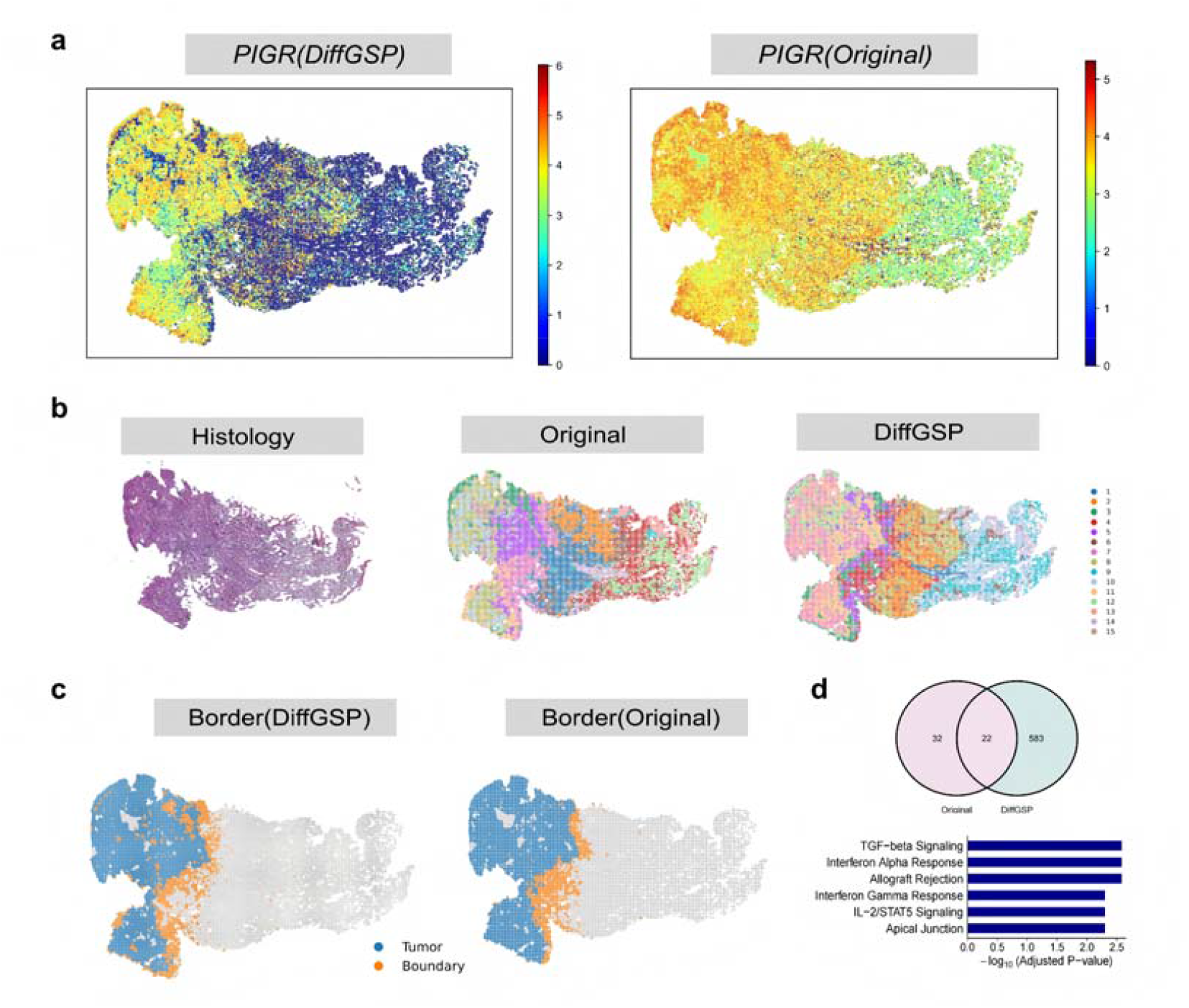
Analysis of human lung cancer Stereo-seq data of P3 tissues. **a**, Diffusion-reduced gene expression patterns in DiffGSP-processed compared to original data in sample P3. **b**, Histology and clustering comparison between DiffGSP-processed and original data. **c**, Tumor and border regions on the basis of the clustering results. **d**, Overlap of boundary-specific DEGs identified from the original and DiffGSP-processed data and hallmark activities of DEGs uniquely detected by DiffGSP.

## Supplementary Information

### Supplementary Figures

**Supplementary Figure 1 | Total count and gene expression pattern examples show the diffusion and the serious dropout phenomenon in ST data**. The color intensity represents the log-transformed total transcript counts or gene expressions of a gene per spot in mouse brain ST data.

**Supplementary Figure 2 | Total count and gene expression pattern examples show the diffusion phenomenon in Visium data**. The color intensity represents the log-transformed total transcript counts per spot in mouse brain, mouse kidney and human colorectal cancer Visium data. White contours indicate the tissue boundaries.

**Supplementary Figure 3 | Total count and gene expression pattern examples show the diffusion phenomenon in VisiumHD data with different resolutions**. The color intensity represents the log-transformed total transcript counts per spot in human colorectal cancer VisiumHD data. White contours indicate the tissue boundaries.

**Supplementary Figure 4 | Total count and gene expression pattern examples show the diffusion phenomenon in Stereo-seq data**. The color intensity represents the log-transformed total transcript counts per bin (bin50) in mouse lung cancer Stereo-seq data. White contours indicate the tissue boundaries.

**Supplementary Figure 5 | Spatial distribution of transcript counts across human–mouse chimeric datasets S1, S2 and S3. a**, Spatial distribution of transcript counts in dataset S1. The left and right columns display the spatial expression patterns of mouse and human transcripts, respectively, across the Original, SpotClean, and DiffGSP, STAGATE and Sprod datasets. **b**, Spatial distribution of transcript counts in dataset S2, displaying the same comparative layout as in (**a**). **c**, Spatial distribution of transcript counts in dataset S3, displaying the same comparative layout as in (**a**). **d**, Comparison of Log Fold Change (LogFC) values across three datasets (S1, S3, and S2) processed by different methods: Original, SpotClean, DiffGSP, STAGATE and Sprod. For human transcripts, LogFC is calculated as the log-transformed ratio of the mean human transcript count in human spots to that in mouse spots. For mouse transcripts, LogFC is computed as the log-transformed ratio of the mean mouse transcript count in mouse spots to that in human spots. Higher LogFC values indicate better tissue specificity and decontamination accuracy.

**Supplementary Figure 6 | Performance comparison on simulated datasets across benchmarking methods. a**,**b**, Performance evaluation of five methods across 15 simulated samples based on (**a**) Mean Squared Error (MSE) and (**b**) Pearson Correlation Coefficient (PCC). The methods evaluated include DiffGSP (pink), SpotClean (purple), Original (green), STAGATE (orange), and Sprod (blue). Each boxplot illustrates the distribution of metrics calculated across all simulated genes within the respective sample. **c**, Spatial expression maps of four representative genes (Gene-1 to Gene-4) demonstrating the capacity of each benchmarking method to recover the ground-truth, pre-diffusion spatial patterns.

**Supplementary Figure 7 | Influence of the diffusion coefficient** a **on inverse diffusion reconstruction. a**, Spatial expression recovery maps of a representative gradient gene (Gene A) modeled across a spectrum of diffusion coefficients *a*, ranging from 0.0067 to 0.4956) without automated parameter correction. The dashed box highlights the optimal reconstruction result yielding a stable spatial pattern, where the diffusion coefficient (a = 0.353) is adaptively determined by DiffGSP using the knee-point criterion. Beyond this knee point, the reconstruction undergoes severe numerical instability and boundary artifacts due to the ill-posed nature of the inverse problem.

**Supplementary Figure 8 | Computational efficiency and scalability benchmarking of DiffGSP. a**, Execution time (in seconds) on small-to-medium-scale datasets ranging from 784 to 19,600 spots. **b**, Peak CPU memory usage (in megabytes, MB) across small-to-medium-scale datasets. **c**, Peak GPU memory usage (in MB) for DiffGSP across the small-to-medium-scale datasets. **d**, Execution time on large-scale datasets ranging from 28,224 to 78,400 spots, utilizing the subgraph optimization strategy (DiffGSP-subgraph). **e**, Peak CPU memory usage across the large-scale datasets. **f**, Peak GPU memory usage for DiffGSP-subgraph across the large-scale datasets. Value labels above each bar indicate exact measurements. Zero values for SpotClean in (**d**) and (**e**) denote execution failures due to Out-Of-Memory (OOM) errors. Zero values for SpotClean in (**c**) and (**f**) denote that SpotClean does not occupy GPU resources.

**Supplementary Figure 9 | Hyperparameter sensitivity analysis of DiffGSP**. Performance stability of DiffGSP evaluated under varying combinations of key hyperparameters on simulated datasets. **a**, Heatmaps showing the Pearson Correlation Coefficient (PCC; left) and Mean Squared Error (MSE; right) across joint variations of the inverse diffusion iteration steps *k* (analogous to the reverse time horizon) and the initial diffusion coefficient *a*. **b**, Heatmaps of PCC (left) and MSE (right) across joint variations of the inverse diffusion steps *k* and the network-based spatial smoothing parameter *C*. The high and stable PCC values alongside low MSE scores across a wide range of parameter combinations demonstrate the robustness of DiffGSP. Note that the initial diffusion coefficient *a* is adaptively optimized during execution to prevent the system from entering an ill-posed regime.

**Supplementary Figure 10 | Spatial domain clustering comparison on the high-resolution Stereo-seq mouse brain dataset. a**, Annotation information of the mouse brain hemisphere section from the previous study, serving as the reference for spatial structures. **b**, Unsupervised spatial clustering results generated from the original data. **c**, Unsupervised spatial clustering results generated from the DiffGSP-processed data.

**Supplementary Figure 11 | Spatial expression patterns of characteristic constituent genes for Gene modules identified in DiffGSP-processed data. a**, Spatial distribution map of the module eigengenes for Gene Module 2 (GM 2) (left), accompanied by the spatial expression maps of six representative associated genes (*Aldob, Fth1, Galnt11, Pdzk1, Prdx5*, and *Pth1r*) across the mouse kidney section (right). **b**, Spatial distribution map of the module eigengenes for Gene Module 6 (GM 6) (left), along with the spatial expression maps of six representative associated genes (*Cyp2d26, Igfbp4, Miox, Slc4a4, Slc5a2*, and *Slc22a8*) (right).

**Supplementary Figure 12 | Spatial domain analysis of transcriptional and functional heterogeneity within tumor subregions. a**, Local enlarged view of spatial domains of three ROIs. **b**, Differential expression heat maps of tumor regions with spatial heterogeneity. **c**, Hallmark activities of DEGs in ROI 1-Tumor I and Tumor II.

**Supplementary Figure 13 | Analysis of human colorectal cancer VisiumHD data of P2 tissues with 16um and 8um resolution. a**, Comparison of cell type identification between original and DiffGSP-processed data in human colorectal cancer VisiumHD data of P2 tissue with 16um and 8um resolution. **b**, Tumor and border regions. Tumor region is defined as the region labeled with tumor cells. Tumor border is defined as the spots within 6.25 radius of the tumor region expanding outward, and some isolated spots are removed. **c**, Comparison of the expression of tumor marker genes increased in the tumor area. Tumor marker genes were selected by differential expression analysis of matched scRNA-seq. And we filtered out spots expressed as zero in tumor in the original data.

**Supplementary Figure 14 | DiffGSP-enhanced spatial gene expression patterns and transcriptional heterogeneity within tumor border subregions. a**, Diffusion-reduced gene expression patterns in DiffGSP-processed data, compared to original data in sample P1. **b**, Cell type proportion of tumor, mixed border, and fibroblast-domain in original data. **c**, Diffusion-reduced gene expression patterns in DiffGSP-processed compared to original data in tissue sample P4. **d**, Volcano plot of DEGs between Border I and Border II from the DiffGSP-processed dataset.

